# A new phylogeny and phylogenetic classification for Solanaceae

**DOI:** 10.1101/2025.07.10.663745

**Authors:** Rocío Deanna, Gloria E. Barboza, Lynn Bohs, Steven Dodsworth, Edeline Gagnon, Leandro L. Giacomin, Sandra Knapp, Andrés Orejuela, Peter Poczai, Tiina Särkinen, Stacey D. Smith, Richard G. Olmstead

**Author notes:** Author for correspondence: Richard G. Olmstead.

## Abstract

Solanaceae are a globally distributed family of considerable economic importance. Recent phylogenetic work on various clades in the family has resulted in a plethora of names at various taxonomic levels. This results in a confusing set of names applied to clades at different ranks, and to the use of informal names that are often not accepted in online resources. To solve this problem, we present here a phylogenetic classification following the *PhyloCode* for the Solanaceae. Traditional taxon names previously recognized at the ranks of tribe, genus, and section were maintained when associated with clades. We base our clade definitions on a newly constructed phylogeny of the family using 10 plastid and nuclear markers with a broad sampling of 1,474 species, roughly half of the diversity of the family. We complement this marker-based analysis with a phylogenomic analysis of 110 taxa (including 97 of the 102 genera in the family) using the Angiosperms353 probe set. This pair of analyses produced highly concordant phylogenies and provides strong support for the clades included in the classification. In total, we propose and define rank-free names for 38 major clades, including 16 infrageneric taxa within the mega-diverse *Solanum*. This rank-free system intends to promote name stability as new data become available, avoiding name changes to well-known groups.

## INTRODUCTION

Solanaceae are a plant family with approximately 3,000 currently accepted species, almost half of them in the mega-diverse genus *Solanum* L. (World Flora Online Consortium, 2024). The family has long been considered coherent; early herbalists (e.g., Gerard, 1597) recognized affinities among these plants, mostly based on medical usage and overall morphological similarity. Linnaeus (1753) grouped most of the currently recognized genera that he knew together in his “Pentandria Monogynia” (five stamens, one ovary), and A. de Jussieu (1789) first coined the family name, based on the genus *Solanum*, which was and still is the largest genus in the family. Members of the family occur worldwide on all continents except Antarctica, with a center of generic and species diversity in South America (Olmstead, 2013; Dupin & al., 2016). Species of Solanaceae are found in a huge range of habitats, from dry deserts in Australia and western South America to hyper-wet cloud forests in the Andes and Himalayas. Habit is similarly diverse in the family, ranging from annual herbs to tall trees reaching 30 m in height, and trait variation is also diverse (Knapp, 2002a, 2010; Zhang & al., 2017). The last treatment of the family at a global level with complete species descriptions was Dunal (1852).

Since the explosive increase in knowledge of Solanaceae that occurred with the collections of botanists and plant collectors like Alexander von Humboldt, Aimé Bonpland, and Karl von Martius in the early part of the 19th Century (Knapp, 2007; Sobral & Stehmann, 2009; BFG, 2022), the family has expanded in diversity. Current digitization of herbaria has made this diversity even more accessible (Versiane & al., 2025). Dunal’s treatment of the family (Dunal, 1852) recognized two tribes (Nolaneae and Solaneae, based on the number of locules in the ovary), with the Solaneae comprising nine subtribes. Traditional classifications of the family in the late 20th Century generally recognized infrafamilial divisions at the rank of subfamily, ranging from two (Cestroideae and Solanoideae; D’Arcy, 1979; Hunziker, 1979) to six (Cestroideae, Solanoideae, Anthocercidoideae, Juanulloideae, Salpiglossoideae, and Schizanthoideae; Hunziker, 2001). Several genera, however, have been variously excluded or included in different systems due to their seemingly aberrant morphology. The family Nolanaceae was erected by Dumortier (1829) to accommodate the genus *Nolana* L., the herbaceous genera *Schizanthus* Ruiz & Pav. and *Salpiglossis* Ruiz & Pav. were placed in the Scrophulariaceae by Bentham (1846), the canopy tree *Duckeodendron* Kuhlm. was initially placed in Solanaceae (Kuhlmann, 1925) but later segregated at the family level (Kuhlmann, 1930), *Sclerophylax* Speg. was recognized at the family level by Hunziker (2001) and the Antillean genera related to *Goetzea* Wydl. were excluded from the family by Hunziker (2001). Modern phylogenetic studies using DNA sequence data have shown that all of these genera previously recognized as distinct families allied to the Solanaceae are nested within a monophyletic Solanaceae (Fay & al., 1998; Olmstead & al., 2008; Särkinen & al., 2013; Ng & Smith, 2016; this paper).

The broad picture of Solanaceae relationships based on explicit phylogenetic analysis has been understood for over 25 years. From the earliest cpDNA studies (Olmstead & Palmer, 1992; Olmstead & al., 1999, 2008), it was apparent that some of the traditional systematic divisions within the family, including a primary division into the two subfamilies Cestroideae and Solanoideae, were inconsistent with contemporary systematic philosophy that named taxa should be monophyletic. At the same time, other well-supported clades previously unrecognized in conventional classifications of Solanaceae were identified; many of these were given informal names, including the large “X=12” clade of taxa with 12 chromosomes as the base number. Attempts to revise the classification of Solanaceae within the traditional rank-based Linnaean system applying the rules of the *International Code of Botanical Nomenclature* (now *International Code of Nomenclature for algae, fungi, and plants*, or hereafter the *Code*; Turland & al., 2018) have resulted in revised circumscriptions of infrafamilial taxa at all ranks (Olmstead & Bohs, 2007; Reveal, 2012; Barboza & al., 2016), but have remained insufficient to provide stable names for all of the clades identified and used to communicate about diversity in Solanaceae, which, after all, remains the fundamental reason for classification and the rules of nomenclature by which taxa are named.

The *Code* is a series of requirements and recommendations for naming taxa using a hierarchical series of ranks, with certain of these being required, and therefore privileged in usage (e.g., genus, family). While phylogeny is, by its nature, hierarchical, that hierarchy is not uniformly structured in a way that fits neatly into the ranks established by the *Code*. Thus, the need often arises for taxa to be named that do not fit into a rank-based system, or for names to be changed simply to accommodate changes in rank. The *PhyloCode* (Cantino & de Queiroz, 2020) provides a system in which taxa in a classification are clades explicitly defined by reference to phylogeny and given names tied to that definition. Clades may be named at any level in the phylogenetic hierarchy, and since there are no ranks, all are equally privileged.

Advances in genome-scale data gathering methods and their phylogenetic analyses have dramatically expanded sampling of both species and data in phylogenetic studies of Solanaceae since those early studies, and have continued to refine our understanding of the family’s phylogeny (Goldberg & al., 2010; Särkinen & al., 2013; Ng & Smith, 2016; Deanna & al., 2019). By compiling data from the many studies over the past 35 years to include 1474 species of Solanaceae, we provide the broadest phylogenetic study of the family to date. We also leverage high-throughput sequencing methods (Zuntini & al., 2024) to complement this marker-based phylogeny with a genome-scale analysis for 111 taxa with representation of nearly all genera in Solanaceae. Here, we use these phylogenetic trees as the basis for a revised classification of Solanaceae with taxon names explicitly tied to clades by use of clade definitions as laid out in the *PhyloCode* (Cantino & de Queiroz, 2020; http://phylonames.org/code/).

## MATERIALS AND METHODS

### Phylogenetic nomenclature

Following the *PhyloCode* principles (Cantino & de Queiroz, 2020; http://phylonames.org/code/), clades names are given clade definitions, which may be minimum clades (e.g., the smallest clade that contains two or more specifiers) or maximum clades (e.g., the largest clade that includes one specifier, but excludes one or more specifiers). Specifiers are species included in a phylogenetic tree designated as the reference phylogeny for the clade definition. Secondary reference trees may be designated that support or provide additional clarification for the clade being named. In most cases, the primary reference phylogeny is the tree derived from the analysis of the marker-based 1475-taxon dataset (**Fig. 1**, see Figs. S1 and S2 for the full tree).

**Figure 1.**
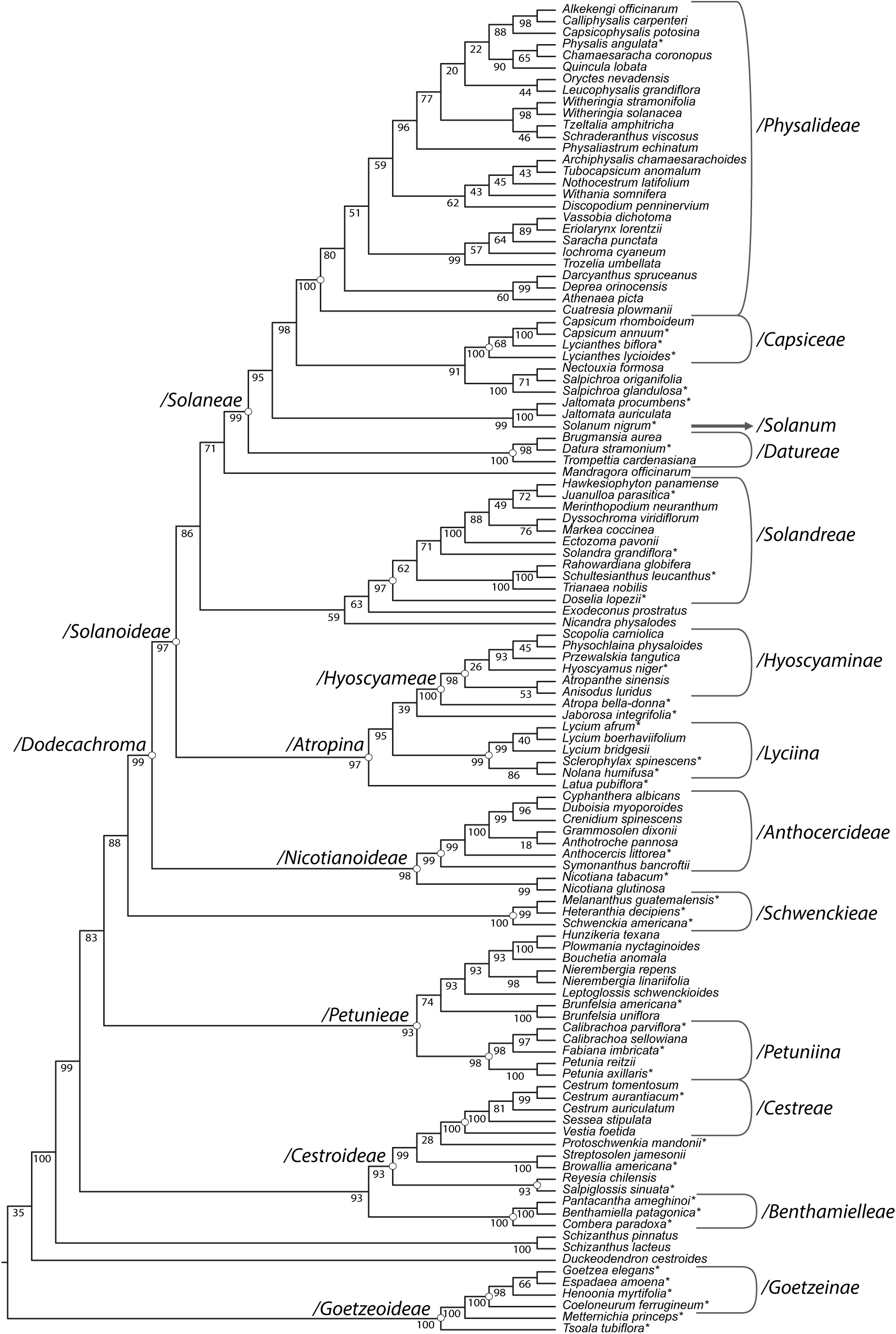
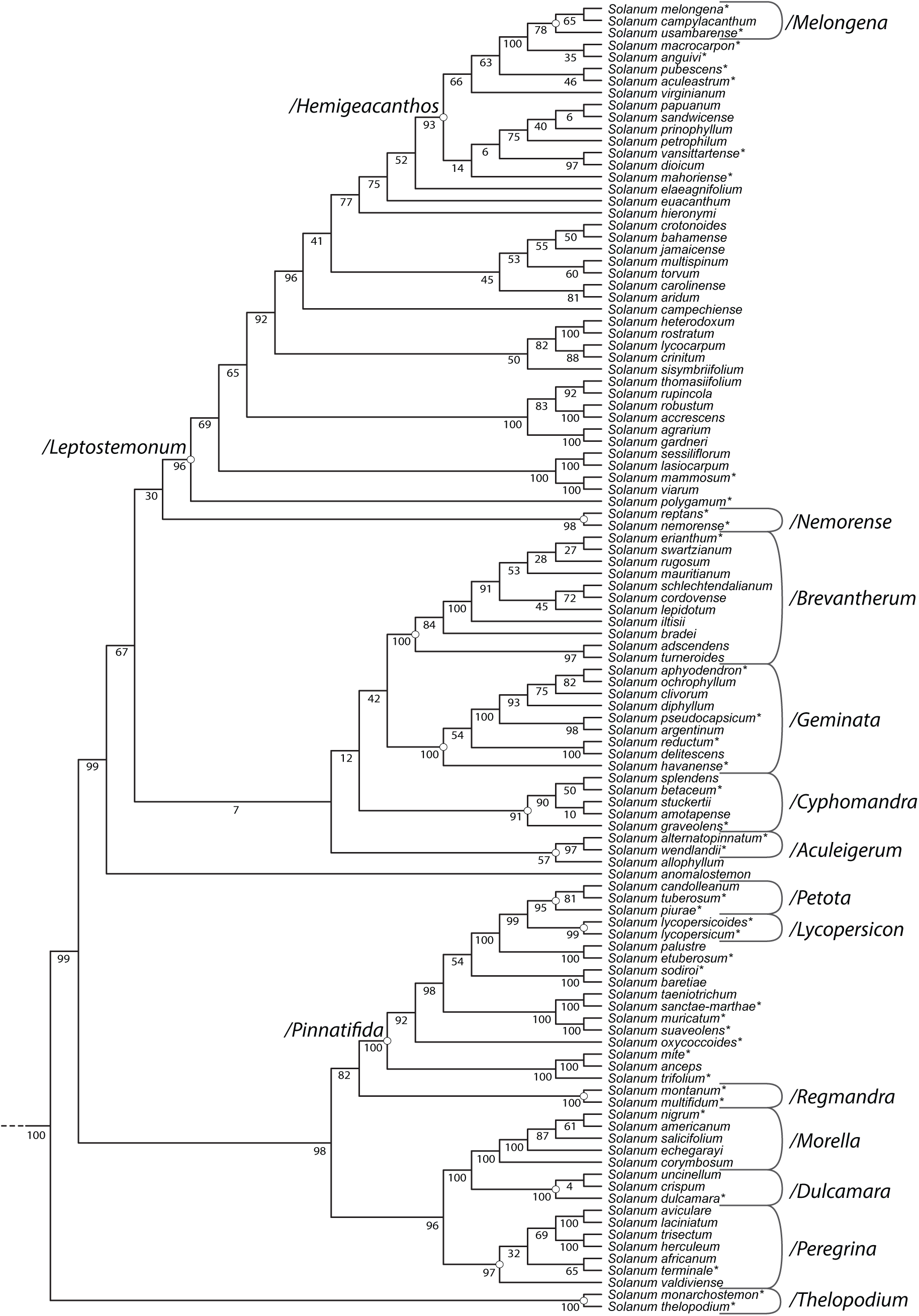
Phylogenetic relationships based on the 10-gene analysis of 1474 Solanaceae species. (A) Family-level phylogeny, pruned from the full tree to include selected representatives of major lineages. Specifiers used in clade definitions are indicated with asterisks. Branches are annotated with bootstrap support values. Named clades are shown either to the left of the node or with brackets on the right. The corresponding node is also marked with an open circle. All clade names are equal in font size to communicate that all are equally privileged. The arrow indicates where the tree for *Solanum* continues in Fig. 1B. (B) Phylogeny for selected *Solanum* species, pruned from the full tree. Annotations follow (A).

### Marker-based phylogenetic inference

We estimated a maximum likelihood phylogeny for 1,474 species of Solanaceae using 10 nuclear and plastid regions: ITS, LEAFY, *matK*, *ndhF*, *ndhF-rpl32*, *psbA*, *rbcL*, *trnLF*, *trnSG*, and *waxy*. We used the PyPHLAWD pipeline (Smith & Walker, 2018) to gather sequence data from these 10 regions from GenBank. We targeted taxa included in two previous marker-based family-level analyses (Särkinen & al., 2013; Ng & Smith, 2016) and added 234 new taxa compared to these studies (Table S1). We aligned sequences using the MUSCLE algorithm (Edgar, 2004) in Mega v. 12 (Kumar & al., 2024). The 10-marker matrix for 1575 taxa was cleaned by pruning rogue taxa (i.e., unstable terminals causing lower branch support) using the software RogueNaRok (Aberer & al., 2013). Two iterations of RogueNaRok were performed based on trees generated from fast bootstrap analyses with RAxML-VI-HPC v.8 (Stamatakis, 2014) in the CIPRES Science Gateway (Miller & al., 2010). For these RAxML analyses, we partitioned the dataset by gene region and used a GTR + Γ model for each. We assessed support with 100 bootstrap replicates. For the RogueNaRok analyses, we applied a 50% bootstrap consensus threshold and a drop set size of one for the identification of rogue taxa. We removed a total of 100 rogue taxa (Table S2), most of them with a large amount of missing information. The final matrix comprised 10,914 bp of aligned sequenced data and 1,474 Solanaceae species, plus the outgroup *Ipomoea purpurea* (L.) Roth (Convolvulaceae). We carried out the final partitioned maximum likelihood search on the combined dataset, again in RaxML v.8 with the GTR + Γ model, and completed 1,000 bootstrap replicates to measure clade support. The final alignment, partitions, and trees are available in the OSF repository (https://osf.io/zem8b/?view_only=91df995fc9e54beeae90f24bbea7dfe1).

### Taxon sampling, DNA extraction, and a353 sequencing

We sampled 97 currently recognized genera of Solanaceae, with the exception of *Darcyanthus* Hunz. ex N.A.Harriman, *Mellissia* Hook. f., *Combera* Sandwith, *Calliphysalis* Whitson, and *Melananthus* Walp. We included multiple species per genus in larger or polyphyletic groups (e.g., *Solanum*, *Lycianthes* Hassl.*, Physalis*), based on prior phylogenetic evidence (Zamora-Tavares & al., 2016; Orejuela & al., 2017; Spalink & al., 2018; Deanna & al., 2019). A total of 110 ingroup species were included, with *Ipomoea purpurea* as the outgroup. Full voucher information is provided in Table S3. Genomic DNA was extracted from ∼2 cm² of leaf tissue using Qiagen DNeasy Plant Mini Kits, with multiple extractions pooled for older herbarium material. DNA concentration and integrity were assessed using a Qubit® 3.0 Fluorometer and 1.0× agarose gel, respectively. High-quality DNA was sheared to ∼350 bp (when needed) using a Covaris M220. Double-indexed Illumina® libraries were prepared with NEBNext® Ultra™ II kits and quantified via Qubit® and TapeStation. Equimolar libraries (1 µg) were pooled and enriched using the Angiosperms353 v1 probe set (Arbor Biosciences). Hybridizations were conducted at 65°C for 28–32 h, followed by PCR amplification (10 cycles) and purification. Libraries were sequenced on an Illumina MiSeq using v2 (300-cycle) and v3 (600-cycle) kits at the Royal Botanic Gardens, Kew.

### Read processing and target assembly

To complement our sampling, additional sequencing data were retrieved from the NCBI Sequence Read Archive (SRA; Table S3). Data acquisition was performed using the SRA Toolkit, and individual datasets were extracted using the fastq-dump utility. Raw reads were first filtered for exogenous DNA using FastQ Screen (Wingett & Andrews, 2018), removing low-complexity sequences with multiple hits across genomes. A second filtering step excluded contaminants typical of herbarium specimens and surface microbiota by aligning reads to a custom Bowtie2 (Langmead & Salzberg, 2012) database derived from genomes listed in Bieker & al. (2020). Adapter sequences and low-quality bases (Phred <20; read length <60 bp) were trimmed using BBDuk from the BBTools package (Bushnell, 2014). Read quality was evaluated with FastQC (Andrews, 2010). A matrix comprising targets from the Angiosperms353 (A353) reference set (Johnson & al., 2019) was used as input for HybPiper v.2.3.2 (Johnson & al., 2016). Raw reads were aligned to the target sequences using BWA v.0.7.17 (Li & Durbin, 2009) and subsequently sorted and indexed with SAMtools v.1.17 (Li & al., 2009). Gene assemblies were conducted with SPAdes v.3.15.5 (Bankevich & al., 2012), adjusting the minimum coverage threshold (--cov_cutoff) to 5 to increase contig retention for low-coverage regions (Liu & al., 2021). Resulting contigs were stitched into full-length gene sequences using Exonerate v.2.4.0 (Slater & Birney, 2005). Recovery efficiency across samples was evaluated using the “hybpiper stats” and “hybpiper recovery_heatmap” functions. All analyses were conducted on the Puhti supercomputing infrastructure at CSC – IT Center for Science, Espoo, Finland.

### Phylogenomic analyses

Assembled sequences were processed using custom scripts (<u>OSF repository link</u>). Genes flagged with paralog warnings above the threshold were excluded. Remaining genes were aligned individually with MAFFT v7.49 (Katoh & Standley, 2013), allowing up to 30% missing data. Alignments of the final 227 genes were inspected manually and concatenated using BAD2matrix (Salinas & al., 2024). Maximum likelihood (ML) analyses were performed with IQ-TREE v2.3.6 (Minh & al., 2020a). Partitioning was optimized using ModelFinder2 with the -m MF+MERGE option until model fit stabilized (Kalyaanamoorthy & al., 2017; Lanfear & al., 2014). The best ML tree was inferred from 10 independent runs with 1,000 ultrafast bootstrap replicates (Hoang & al., 2018), applying the -bnni option to minimize branch support inflation. Species tree inference followed the multi-species coalescent model in ASTRAL v5.7.8 (Zhang & al., 2018), using gene trees estimated in IQ-TREE v2.3.6 with 1000 bootstrap replicates. Quartet support values (-t 2) were calculated to assess branch reliability. Gene and site concordance factors (gCF and sCF) were estimated in IQ-TREE to evaluate phylogenetic signal beyond bootstrap support (Minh & al., 2020a, 2020b). To investigate the relationship among support metrics, we compared bootstrap values, gCF, and sCF across all internal branches. Following Minh & al. (2020b), we visualized the data using a Viridis-colored scatter plot to highlight instances where high bootstrap support may conceal gene tree discordance or where concordance factors offer additional support. Phylogenetic discordance was assessed by mapping the set of 227 rooted nuclear gene trees onto the species tree using PhyParts (Smith & al., 2015); only bipartitions with bootstrap support ≥50% (-s 0.5) were retained.

## RESULTS

### Marker-based family tree

Our analysis of 1475 taxa using 10 plastid and nuclear markers recovered well-supported relationships for major clades reported in previous family-wide studies (**Fig. 1**, **Fig. S1**). These include the large X=12 clade (BS=88%) and many lineages previously treated as subfamilies (e.g., /*Cestroideae*, /*Goetzeoideae*, /*Nicotianoideae*, and /*Solanoideae*, BS 93-100, **Fig. 1A**). Some clades appear better supported than in previous analyses (e.g., BS=93% for */Petunieae* here vs BS>80% in Särkinen & al., 2013), likely due to the addition of three additional markers and over 200 new tips, even compared to the most recent marker-based analysis at this scale (Ng & Smith, 2016). We also note that the small Chilean genus *Reyesia* Clos, which has varied in position across past studies, is now clearly associated with the morphologically similar *Salpiglossis* Ruiz & Pav. (**Fig. 1A**).

Nevertheless, the position of the root remains unstable in this analysis, as in past studies. Here, the maximum likelihood tree places */Goetzeoideae* as sister to the rest of the family, albeit with low support (BS=35%). Other clades have been recovered as possible sister groups, e.g., *Schizanthus* (Olmstead & al., 2008; Goldberg & al., 2010; Huang & al., 2023), *Schizanthus* + /*Goetzeoideae* + *Reyesia* + *Duckeodendron* (Särkinen & al., 2013), *Schizanthus* + /*Goetzeoideae* (Ng & Smith, 2016), all of which used the same outgroup taxa from Montiniaceae and Convolvulaceae. Better resolving these early divergences will be particularly important for making inferences about the biogeographic history, since /*Goetzeoideae* occur in Madagascar (*Tsoala*) and the Antilles (*/Goetzeinae*), whereas *Schizanthus* only occurs in southern South America.

Our phylogenetic results for *Solanum* mirror prior analyses using similar sets of markers, with some major and minor clades strongly supported but others having short branches and low bootstrap supports. For example, the large clade */Leptostemonum* comprising the ca. 450 “spiny solanums” is recovered with 96% bootstrap support (**Figs. 1B**, S1), consistent with its monophyly in early studies (e.g., Bohs & Olmstead, 1997; Olmstead & Palmer, 1997; Levin & al., 2006). Nevertheless, many splits within this clade are largely unresolved (see also Gagnon & al., 2022). The short internal branches in many portions of the *Solanum* phylogeny (Fig. S2) have been largely attributed to incomplete lineage sorting associated with rapid speciation (Gagnon & al., 2022), especially in /*Hemigeacanthos* (Echeverría-Londoño & al., 2020).

### A353 Gene Recovery

Recovery statistics varied considerably across taxa, reflecting differences in input read depth, phylogenetic distance from reference taxa, and library quality (Table S4). On average, over 300 genes were mapped per taxon, with many species achieving high completeness scores: several taxa recovered over 250 genes at ≥50% length and more than 100 genes at ≥75% target length. The average on-target rate across samples ranged between 1– 8%, consistent with expectations for hybrid enrichment in a large, phylogenetically diverse clade. The samples with lower recovery were typically characterized by low total read counts, reduced on-target percentages (often below 2%), and limited recovery of stitched contigs. Additionally, although paralog warnings were generally low across samples, some low-performing samples recovered high paralog warnings or showed a high number of genes without assembled contigs. Nevertheless, the recovery heatmap (Fig. S3) reveals a broad consistency in gene recovery across loci, with a core set of high-confidence targets captured in nearly all taxa, providing a basis for phylogenetic inference.

### Phylogenomic analyses using A353 data

The concatenated maximum likelihood and species tree analyses of the A353 data revealed remarkable congruence with the marker-based tree, with all of the clades included in the *Phylocode* classification presented below recovered in both trees (**Figs. 2**, S4). This level of agreement between the 10-marker dataset and the genomic dataset is perhaps unsurprising given that our classification focuses on well-established clades that have been consistently estimated across prior studies.

**Figure 2.**
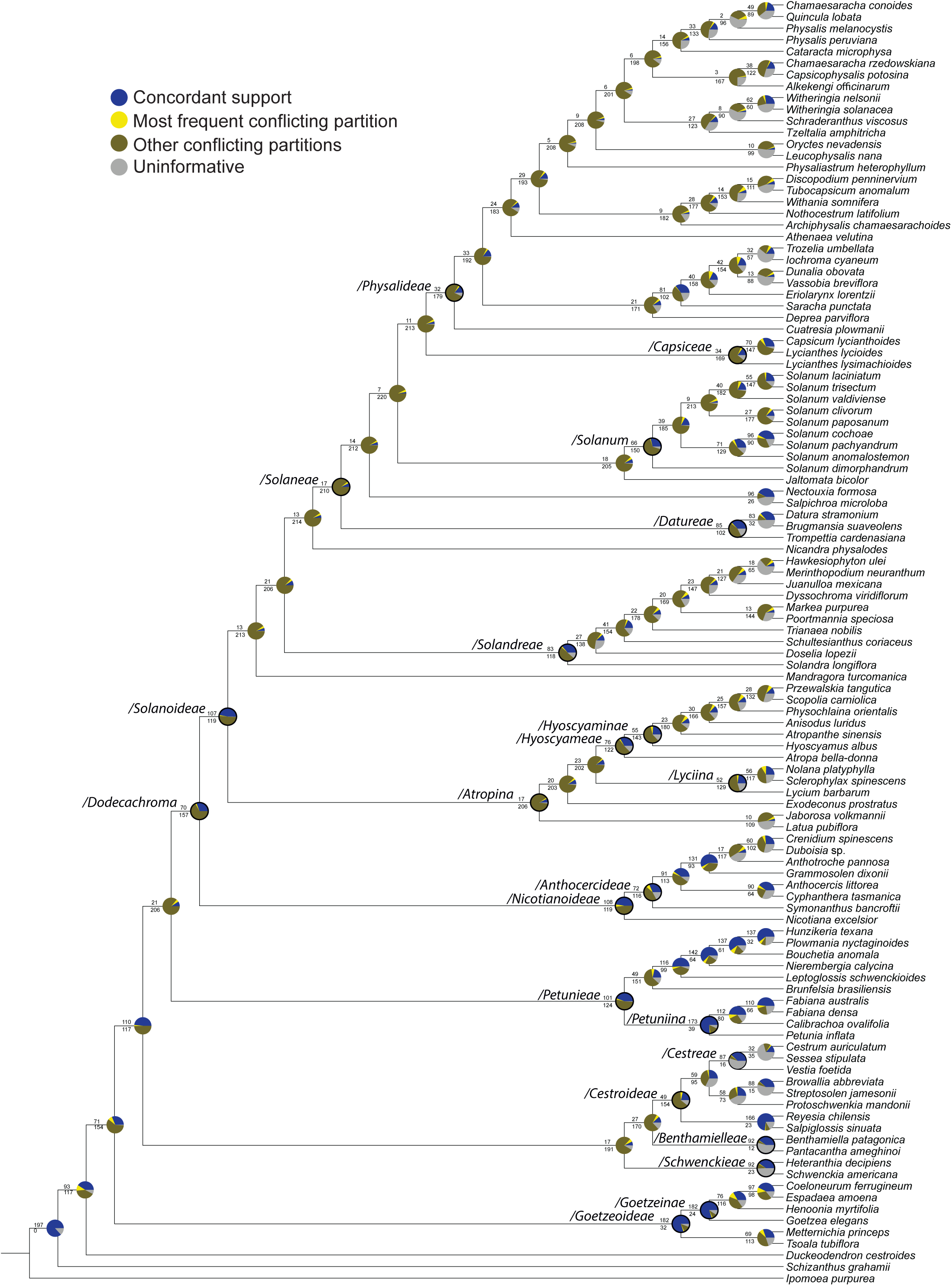
Gene tree conflict and concordance across the species tree based on Angiosperms353 loci. Results are shown as a pie chart on each internal node of the phylogeny. Each pie chart indicates the proportion of gene trees supporting the species tree topology (blue), the most frequent conflicting bipartition (yellow), other conflicting bipartitions (brown), and uninformative gene trees with less than 50% bootstrap support at that node (gray). Clade names follow Fig. 1. Pie charts for named nodes are indicated with a bold circle.

Despite this overall agreement across analyses and datasets, our assessments of concordance among gene trees indicate that the species tree is commonly supported by only a fraction of the genome. At many nodes across the phylogeny, a majority of the genes present a topology different from the species tree with moderate bootstrap support (BS>50%) and another portion are uninformative (i.e., less than 50% for any topology; **Fig. 2**). Moreover, many of the most discordant nodes, both in terms of site and gene concordance factors, are those with very high bootstrap support (>95%, Fig. S5). Thus, the high bootstrap values across the concatenated ML trees likely derive from the (sometimes small) portions of the genome that are both informative and concordant with each other and with the species tree. This pattern of high bootstrap support in concatenated analyses, even with substantial genome-wide discordance, is common in phylogenomic analyses (e.g., Gagnon & al., 2022; Pezzi & al., 2024) and can arise from many factors, including incomplete lineage sorting, hybridization, and polyploidization (Degnan & Rosenberg, 2009; Ning & al., 2024).

### Clade names in Solanaceae

A total of 38 clades are defined (25 minimum clades and 13 maximum clades), including 21 suprageneric taxa in conventional rank-based classification, one at the rank of genus (*Solanum*), and 16 infrageneric taxa within *Solanum* (**Figs. 1-2**; Table 1; Appendix 1). Thirty-three are converted clade names, of which 30 are validly published under the *Code,* and three names were informally assigned to clades in previous phylogenetic studies, and five are first used here. All clade names under *PhyloCode* are written in the Latin alphabet. A slash mark (/) is used simply to denote a formally defined clade name when confusion with a name published under the conventional *Code* may otherwise result, but it is not a required part of the name.

**Table 1.**
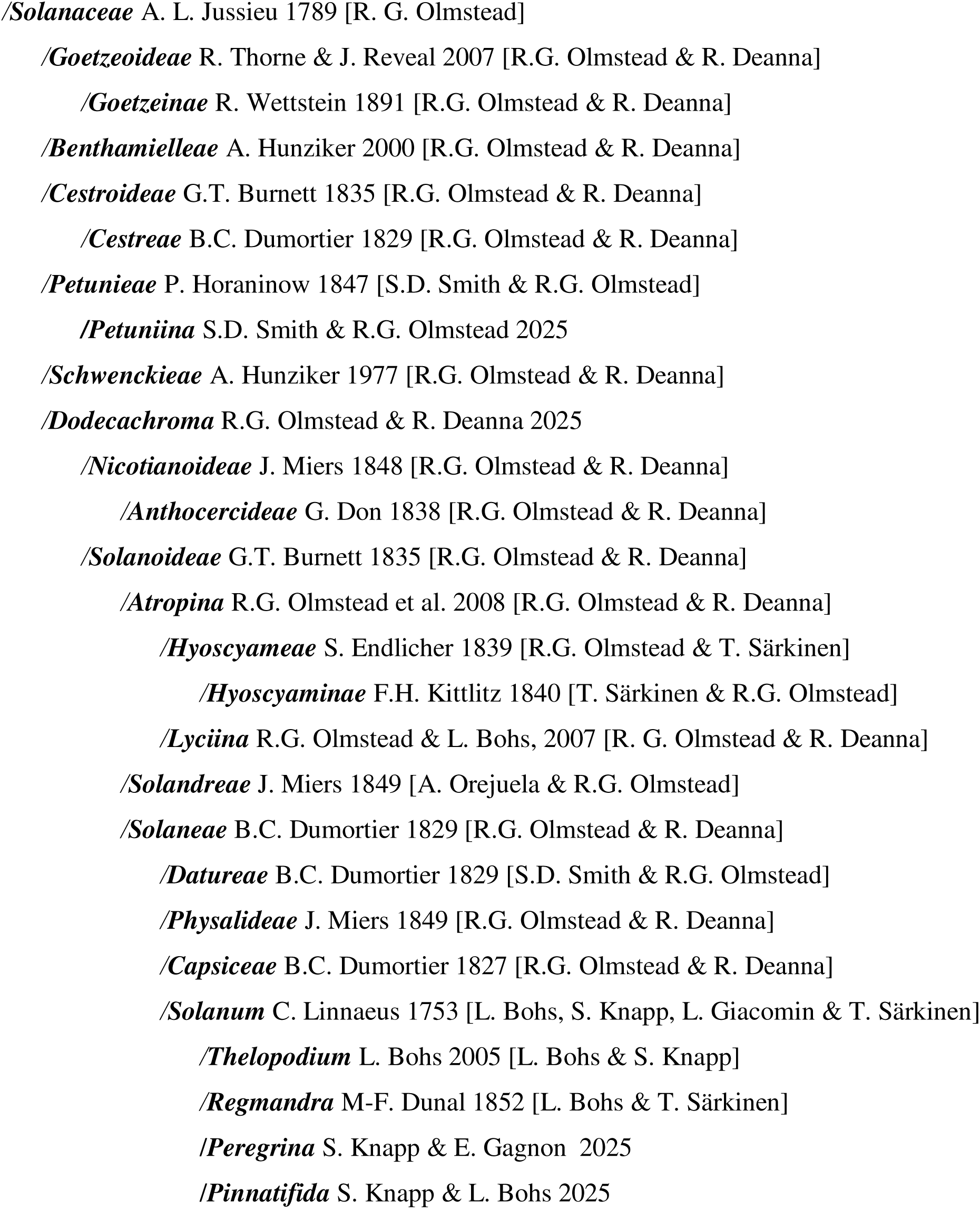
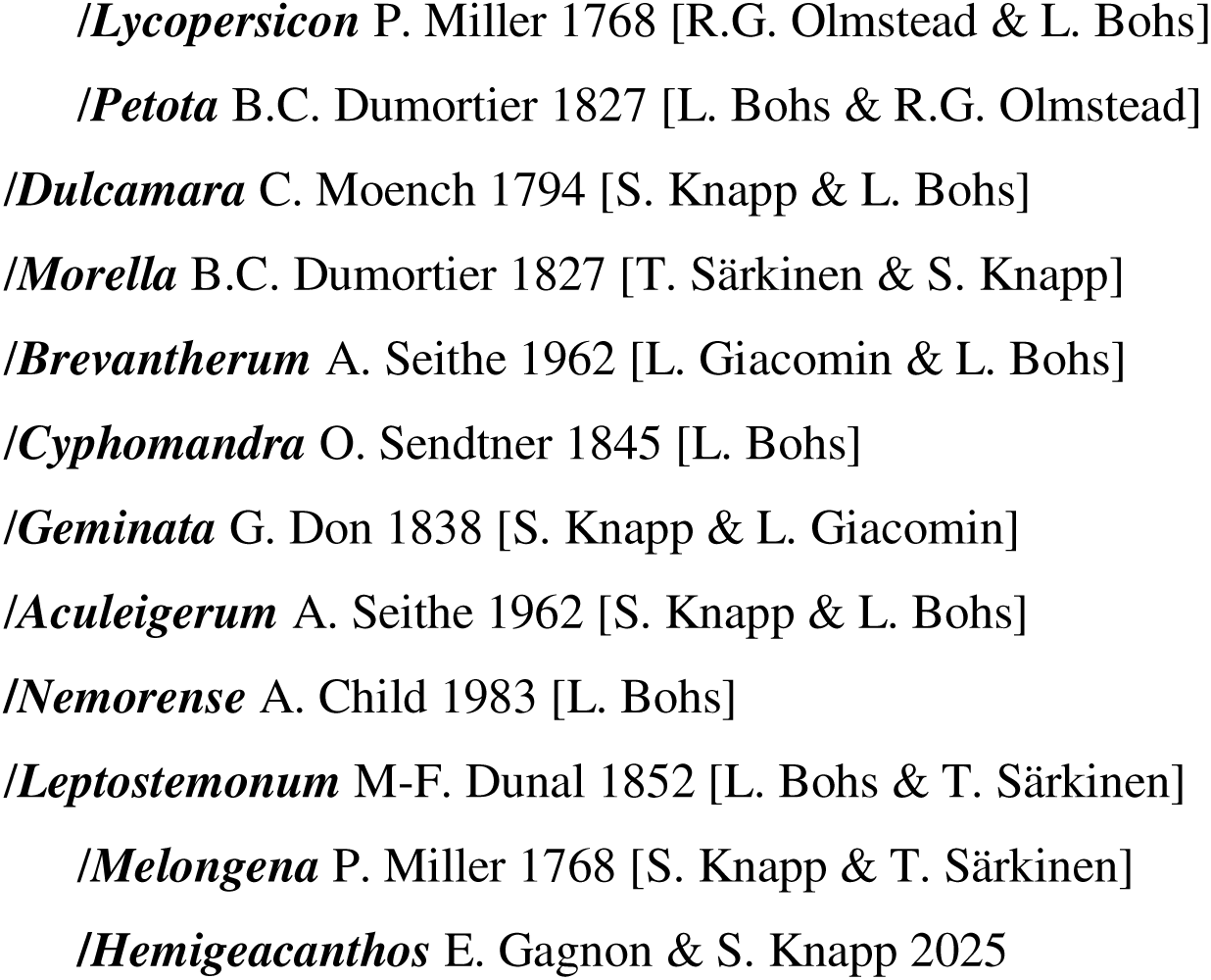
Phylogenetic classification of Solanaceae, including clade name, nominal author and date of first publication for converted clade names, and definitional authors in square brackets (for new clade names, nominal and definitional authors are the same).

## DISCUSSION

### Phylogenetic relationships across Solanaceae

Our molecular phylogenetic results provide a solid foundation for building a stable and informative classification for the family. The 10-gene analysis of 1475 taxa represents the largest phylogeny yet assembled for the family and supports many major clades that were first detected in the earliest plastid-based analyses (Olmstead & Palmer, 1992; Olmstead & al., 1999, 2008). The addition of nuclear markers has strengthened many of these relationships and confirmed the importance of taxonomically important characters (e.g., bud aestivation, chromosome number, fruit type) long used in rank-based classifications (e.g., D’Arcy, 1979; Hunziker, 2001). Moreover, many well-supported clades, in our analysis and in previous studies, lack any known morphological synapomorphy yet appear so consistently across studies (e.g., /*Petuniina*, /*Peregrina*) as to merit recognition with a clade name. We have elected to name 38 clades across the family, based on their stability across studies, while also hoping that others will expand this classification to many additional groups in the future.

Although many aspects of the Solanaceae phylogeny are well understood, our analyses also underscore how much remains to be explored with future studies. For example, the position of the root node varies across datasets and analyses (**Fig. 1A**, **Fig. 2**), although *Schizanthus*, *Duckeodendron* and */Goetzeoideae* appear to be strong contenders for the sister group to the rest of the family. Timetree models that estimate the root as part of the phylogenetic inference (Heath & al., 2014) present a viable option for resolving this enigma, but will rely on significant expansion of the Solanaceae fossil record (Deanna & al., 2023). We also face a challenge in increasing taxon sampling of extant species, since some clades are nearly completely sampled (e.g., 80% of Datureae) while others are very poorly represented (e.g., 19% of Cestreae). Knowing that many of these groups are likely to carry histories including episodes of hybridization, continued work to build the phylogeny will benefit from genome-wide approaches that can provide robust estimates of the species tree while also identifying areas of significant discordance.

### Phylogenetic classification and clade definitions

With expanded taxon and DNA sequence sampling for phylogenetic studies in /*Solanaceae* presented here (**Figs. 1-2**, **S1**) and in other recent studies (Olmstead & Palmer, 1992: Yuan & al., 2006; Olmstead & al., 2008; Poczai & Hyvönen 2013; Särkinen & al., 2013; Sanchez-Puerta & Abbona, 2014; Särkinen & al., 2015; Ng & Smith, 2016; Tepe & al., 2016; Orejuela & al., 2017; Aubriot & al., 2018; Dupin & Smith, 2018; Deanna & al., 2019; Tu & al., 2020; Alaria & al., 2022; Chiarini & al., 2022; Gagnon & al., 2022; Yan & al., 2022; Bozan & al., 2023; Huang & al., 2023; Messeder & al., 2024; Pezzi & al., 2024; Maldonado & al., 2025; Sandoval-Padilla & al., 2025), explicit definitions of clades using principles of phylogenetic nomenclature can be established with confidence throughout /*Solanaceae*. Here we formally define many of the suprageneric clades in /*Solanaceae* and infrageneric clades in /*Solanum* recognized both in traditional classifications and informally designated in previous phylogenetic studies (**Appendix 1**). A formal clade definition for /*Solanaceae* was published in *Phylonyms* (Olmstead, 2020), the companion volume published simultaneously with the *PhyloCode*, which represents the starting point for registration of clade names as required under that code.

In implementing the *PhyloCode* in Solanaceae nomenclature, we have used two types of clade-based definitions (see Cantino & de Queiroz, 2020). Minimum clades are defined as the smallest clade containing two or more species, called ‘specifiers,’ as represented on a reference phylogeny. Minimum clades can stand alone without reference to any other branches in a phylogeny and are best applied when there is confidence about content and branching within the clade being named. Maximum clades are defined as the largest clade containing one species, but excluding one or more other species. Maximum clades require resolution among branches adjacent to the clade being named, and are best applied when there is uncertainty regarding relationships within the clade, or there is insufficient sampling to be confident that a minimum clade definition will capture all of the expected clade content within its definition.

For this paper, we elected to focus on suprageneric clades within /*Solanaceae* and clades within /*Solanum*, because there are substantial differences among traditional classifications at these levels and we hope to provide a consensus taxonomy for systematists working on the family in the future and a better and more stable means of communication about groups. Within /*Solanaceae* broadly, we define most of the suprageneric clades recognized in prior studies, except for the clades within /*Physalidae*, where this and prior studies still lack strong support for some internal clades, which should be resolved in a forthcoming study (R. Deanna, personal communication). Within /*Solanum*, we focus on major clades with strong support both here, in Gagnon & al. (2022) and Messeder & al. (2024), leaving many other smaller clades undefined for the time being, but naming those well-supported clades that contain species of economic importance. Several clade names are newly proposed here, both in /*Solanaceae* and within /*Solanum*, where new evidence now permits confidence in their existence as a clade (/*Petuniina* and /*Lyciina*), or where informal vernacular names in wide use are now formalized according to the *PhyloCode* and provide for future nomenclatural stability (/*Dodecachroma*, /*Dulcamara*, /*Morella*, /*Pinnatifida*, /*Hemigeacanthos*, and /*Peregrina*). The phylogenetic classification presented here complements traditional nomenclature without introducing additional ranks or the need to rename groups to accommodate changes in rank.

Taxonomic sampling in the studies cited above and in more focused studies on tribes and genera also permits formal definitions for many clades recognized at the rank of genus or within genera other than *Solanum*, but where there is less confusion among named groups there is less urgency in their establishment and will be the subject of further refinements to the phylogenetic classification of Solanaceae. Ongoing studies of some groups (e.g., tribes Capsiceae and Physalideae) need to be completed before clades within them can be defined with confidence.

## Author contributions

The initial idea to complete a phylogenetic classification of the Solanaceae came from RGO and SK; RD, SDS, AO, PP, RGO, LB, LLG, TS, EG, and SD produced sequences; RD, SDS, AO, PP and SD built the trees; SDS, RD, and RGO prepared the figures; all authors contributed to writing the initial manuscript; clade definitions for Solanaceae were written by RGO, RD, SDS, GB, and AO, for *Solanum* by LB, TS, SK, LLG, and EG; all authors read and edited the final manuscript.

## Acknowledgments

We thank the curators of herbaria worldwide for their care and attention to the preservation of specimens, as well as their willingness to allow their use in downstream research. The CSC – IT Center for Science, Finland, provided free access to high-performance computing resources essential for conducting this research in support of academic study and education. Samples were kindly provided by our generous colleagues P. Gonzales, C.I. Orozco, J.R. Stehmann, O. Vargas-Ponce, G. van der Weerden, and P. Zamora Tavares. We thank R. Cowan for molecular lab support in generating the a353 data.

Funding for this study came from many sources over many years: The National Science Foundation Planetary Biodiversity Inventory program (PBI *Solanum*: a worldwide treatment, DEB-1354791 to SK & LB) and the National Geographic Northern Europe (GEF-NE 49-12 to TS) in part supported sample collection and early sequence generation; LLG is funded by Conselho Nacional de Desenvolvimento Científico e Tecnológico/CNPq (grants 422191/2021-3, 408914/2023-8 and 404996/2024-8); SD and AO were supported by the PAFTOL project (funded by the Calleva Foundation to the Royal Botanic Gardens, Kew); phylogenetic analyses were supported by the National Science Foundation (DEB 1902797 to SDS) and the European Union under the Marie Skłodowska-Curie grant (MSCA agreement No 101151612 to RD).

## Supplementary figures

**Figure S1**. All-compatible bootstrap consensus tree of 1474 Solanaceae species. The topology was inferred through maximum likelihood analysis of 10 markers; bootstrap scores are given for each branch. The tree is rooted on the outgroup (*Ipomoea batatas*). Color coding of taxon names indicate the following: yellow – type species of genera also used as specifiers in clade definitions; blue – type species of genera not used as specifiers in clade definitions; green – specifiers used in clade definitions that are not type species; pink – placeholder taxa representing closest relatives included in the tree for type species that are not included.

**Figure S2**. Maximum likelihood phylogeny of 1474 Solanaceae species. Branch lengths are shown in substitutions per site.

**Figure S3.** Heatmap showing locus recovery for the Angiosperms353 (a353) target capture set across all sampled taxa. Each row represents a sample, and each column corresponds to a target locus. Color intensity indicates the proportion of each locus successfully recovered by HybPiper, with darker shades representing higher recovery.

**Figure S4.** Maximum-likelihood phylogenetic tree of *Solanaceae* inferred from the Angiosperms353 nuclear gene dataset. Each internal node is annotated with three support metrics: (1) bootstrap support value (BS), (2) gene concordance factor (gCF), and (3) site concordance factor (sCF), in the format BS/gCF/sCF. Bootstrap values were computed with ultrafast bootstrapping; gCF and sCF were calculated using IQ-TREE. The high-resolution support values help illustrate varying levels of phylogenetic confidence and concordance across the tree.

**Figure S5.** Scatter plot visualizing the gene concordance factor (gCF) on the x-axis and the site concordance factor (sCF) on the y-axis, with bootstrap support color-coded using a Viridis color scale. The diagonal line represents the theoretical scenario where gCF and sCF values are equal. Points falling on this line indicate branches where both gene trees and site patterns show equal support.

## Supplementary tables

**Table S1.** Taxa and corresponding references for species added to the dataset after Särkinen & al. (2013). This table lists newly included taxa along with the sources supporting their phylogenetic placement or taxonomic identification.

**Table S2.** Rogue taxa removed after two rounds of RogueNaRok cleaning (Aberer & al., 2013) based on trees generated from fast RAxML bootstrap analyses, using RAxML-VI-HPC v.8 (Stamatakis, 2014). Where the name in the original dataset is the same as the accepted name, the second column is left blank.

**Table S3.** Voucher information for all samples included in the Angiosperms353 (a353) dataset. Includes species names, collector and collection number, herbarium acronym, and accession codes if the data was acquired from the NCBI Sequence Read Archive.

**Table S4.** Summary statistics of HybPiper recovery for the Angiosperms353 (a353) target loci across all sampled taxa. The summary statistics include a comprehensive set of recovery metrics for each sample, such as the total number of loci with sequence data successfully retrieved, the cumulative length of sequences recovered across all loci (in base pairs), the average locus recovery, and the number of loci flagged as putative paralogs, providing insights into capture efficiency and assembly success for each sample.

## Appendix 1. Solanaceae clade names and definitions

> ***/Goetzeoideae*** Thorne & Reveal 2007 [R.G. Olmstead & R. Deanna] converted clade name. **Regnum number**: 1104.—**Definition:** The minimum crown clade containing *Goetzea elegans* Wydler 1830, *Metternichia principis* J.C. Mikan 1823, and *Tsoala pubiflora* Bosser & D’Arcy 1992.

**Primary reference phylogeny:** This paper (Fig. 1A). See also this paper (Figs. 2, S1-S2); Olmstead & al. (2008: Fig. 1). **Etymology**: From genus *Goetzea* Wydler. named to honor the German theologian and zoologist Rev. Johann August Ephraim Goetze, best known for his discovery of tardigrades; the genus was originally described as a member of the Ebenaceae. **Composition:** Eight species currently recognized in the genera *Coeloneurum* Radkl.*, Espadaea* A.Rich.*, Goetzea, Henoonia* Griseb.*, Metternichia* J.C.Mikan, and *Tsoala* Bosser & D’Arcy (Olmstead & al., 2008). **Diagnostic apomorphy**: tricolpate pollen with echinate exine sculpturing and a perforate tectum. **Synonyms:** None. **Comments:** The Antillean genera *Espadaea, Goetzea,* and *Henoonia* were placed in Solanaceae by Wettstein (1891), but subsequently considered to be a distinct family (e.g., D’Arcy, 1991; Hunziker, 2001) following the earlier treatment by Miers (1871), before molecular phylogenetic studies confirmed their place in Solanaceae in a clade with the Brazilian genus *Metternichia* (Fay & al., 1998, Olmstead & al., 1999; Santiago-Valentin & Olmstead, 2003; de Souza & al., 2023) and the monotypic Malagasy genus *Tsoala* (Olmstead & al., 2008). Once it was clear that this clade belonged in Solanaceae, Goetzeoideae came into informal use and was eventually validated by Thorne & Reveal (2007). We choose this name here over the previously used Goetzeaceae, because the composition includes two additional genera (*Metternichia* and *Tsoala*) and to avoid confusion over using a name also recognized at the rank of family under the *International Code of nomenclature for algae, fungi, and plants* (Turland & al., 2018).

> ***/Goetzeinae*** Wettstein 1891 [R.G. Olmstead & R. Deanna] converted clade name. **Regnum number**: 1105.—**Definition:** The minimum crown clade containing *Goetzea elegans* Wydler 1830, *Coelonerurum ferrugineum* (Sprengel) Urban 1899*, Espadaea amoena* A. Richard 1850, and *Henoonia myrtifolia* Grisebach 1866.

**Primary reference phylogeny:** This paper (Fig. 1A). See also this paper (Figs. 2, S1-S2); Olmstead & al. (2008: Fig. 1); Santiago-Valentin & Olmstead (2003: Fig. 1). **Etymology**: From genus *Goetzea* named to honor the Rev. Johann August Ephraim Goetze. **Composition:** Five species currently recognized in the genera *Coeloneurum, Espadaea, Goetzea,* and *Henoonia* (Olmstead & al., 2008). **Diagnostic apomorphy**: scarce endosperm in seeds of ripe fruits. **Synonyms:** Goetzeaceae, Miers 1870. **Comments:** These four Antillean genera have alternately been placed in Solanaceae (e.g., as subtribe Goetzeinae by Wettstein, 1891) or in their own family, Goetzeaceae (Miers, 1871; D’Arcy, 1991; Hunziker, 2001) before molecular phylogenetic studies confirmed their place in Solanaceae (Santiago-Valentin & Olmstead, 2003). The composition of this clade matches that of both the family Goetzeaceae and subtribe Goetzeinae; the latter is chosen for this clade (no name is available at the rank of tribe under the traditional *Code*). Naming a specifier from each genus is necessary due to conflict in resolution of relationships among genera (e.g., this paper, Fig. 2).

> ***/Benthamielleae*** Hunziker 2000 [R.G. Olmstead & R. Deanna] converted clade name. **Regnum number**: 1106.—**Definition:** The minimum crown clade containing *Benthamiella patagonica* Spegazzini 1883, *Combera paradoxa* Sandwith 1936, and *Pantacantha ameghinoi* Spegazzini 1902.

**Primary reference phylogeny:** This paper (Fig. 1A). See also this paper (Figs. 2, S1-S2); Olmstead & al. (2008: Fig. 1). **Etymology**: From genus *Benthamiella* Speg. named to honor George Bentham, 19^th^ century English botanist. **Composition:** Fifteen species in the genera *Benthamiella*, *Combera* Sandwith, and *Pantacantha* Speg. (Olmstead & al., 2008). **Diagnostic apomorphy**: The three genera share a distinctive pollen morphology with large, irregularly shaped exine ornamentations (Stafford & Knapp, 2006), and flowers usually with two opposed bracteoles similar to leaves (except *Pantacantha*; Barboza, 2013). **Synonyms:** None. **Comments:** These three endemic Patagonian genera were placed in either tribes Cestreae or Nicotianeae before molecular phylogenetic studies confirmed the suggestion of Hunziker (2000) to place them in their own taxon, which is resolved as sister to Cestroideae with modest support (Olmstead & al., 2008; this paper, Fig. 1).

> ***/Cestroideae*** Burnett 1835 [R.G. Olmstead & R. Deanna] converted clade name. **Regnum number**: 1107.—**Definition:** The minimum crown clade containing *Cestrum nocturnum* Linnaeus 1753 (*Cestrum aurantiacum* Lindley 1844), *Salpiglossis sinuata* Ruiz & Pavón 1794, and *Browallia americana* Linnaeus 1753.

**Primary reference phylogeny:** This paper (Fig. 1A). See also this paper (Figs. 2, S1-S2); Olmstead & al. (2008: Fig. 1); Montero-Castro & al. (2006: Fig. 1). **Etymology**: From the genus *Cestrum* L., possibly from the Greek “késtron” (sharpness). **Composition:** Clade */Cestreae,* and the genera *Browallia* L., Soler., *Reyesia* Clos, *Salpiglossis* Ruiz & Pav., and *Streptosolen* Miers. **Diagnostic apomorphy**: Pollen having 5-8 colpi and a coarsely striated exine (Gentry, 1979; Stafford & Knapp, 2006). **Synonyms:** None. **Comments:** In most pre-molecular classifications, Solanaceae was divided into two subfamilies, Cestroideae and Solanoideae based on a combination of traits, including fruit type and base chromosome number, but molecular phylogenetic studies have found Solanoideae to be a clade nested in a paraphyletic Cestroideae. Since then, a much-reduced subfamily Cestroideae has been recognized (Olmstead & al., 1999), as defined here. The type species of genus *Cestrum* is not included in the reference phylogeny but is included in Olmstead & al. (2008) and Montero-Castro & al. (2006), where it is most closely related to *Cestrum aurantiacum* included in the primary reference phylogeny.

> ***/Cestreae*** Dumortier 1829 [R.G. Olmstead & R. Deanna] converted clade name. **Regnum number**: 1109.—**Definition:** The maximum crown clade containing *Cestrum nocturnum* Linnaeus 1753 (*Cestrum aurantiacum* Lindley 1844), but not *Browallia americana* Linnaeus 1753 and *Protoschwenkia mandonii* Solereder 1898 and *Salpiglossis sinuata* Ruiz & Pavón 1794.

**Primary reference phylogeny:** This paper (Fig. 1). See also this paper (Figs. 2, S1); Olmstead et al. (2008: Fig. 1); Maldonado & al. (submitted to this volume: Suppl. Fig. 1). **Etymology**: From the genus *Cestrum*, possibly from the Greek word “késtron”, meaning “sharpness.” **Composition:** *Cestrum*, *Sessea* Ruiz & Pav., and *Vestia* Willd. **Diagnostic apomorphy**: Absence of the *Arabidopsis*-type telomeres typical of most angiosperms and found in all other Solanaceae examined (Sykorova & al., 2003); karyotype type III, x = 8, 7 m + 1 sm (Maldonado & al., 2025). **Synonyms:** None. **Comments:** Tribe Cestreae has varied in content in traditional treatments, but usually includes the radially symmetric members of Cestroideae that were not assigned to tribe Nicotianeae. Hunziker (2001) restricted tribe Cestreae to the three genera included here. The type species of genus *Cestrum* is not included in the reference phylogeny, but is included in Olmstead & al. (2008) and Montero-Castro & al. (2006), where it is most closely related to *Cestrum aurantiacum* included in the primary reference phylogeny.

> ***/Petunieae*** Horan 1847 [S.D. Smith & R.G. Olmstead] converted clade name. **Regnum number**: 1110.—**Definition:** The minimum crown clade containing *Petunia axillaris* (Lamarck) Britton, Sterns & Poggenburg 1888 and *Brunfelsia americana* Linnaeus 1753.

**Primary reference phylogeny:** This paper (Fig. 1A). See also this paper (Figs. 2, S1-S2); Olmstead & al. (2008: Fig. 1); Pezzi & al., (2024: Fig. 2). **Etymology**: From *Petunia* Juss., named from the archaic French “petun” or Portuguese “petum”, believed derived from the indigenous Guarani word for tobacco. **Composition:** Clade */Petuniina*, and the genera *Bouchetia* Dunal, *Brunfelsia* L., *Hunzikeria* D’Arcy, *Leptoglossis* Benth, *Nierembergia* Ruiz & Pav., and *Plowmania* Hunz. **Diagnostic apomorphy**: Fruit a septicidal (-loculicidal) capsule, and a heterodynamous androecium (Barboza, 2013). **Synonyms:** None. **Comments:** Most genera included here are assigned to the large tribe Nicotianeae in most traditional classifications (e.g., D’Arcy, 1991; Hunziker, 2001). However, molecular phylogenetic studies place the genus *Nicotiana* with other groups having a base chromosome number of X=12 in /*Dodecachroma* and recognize a clade comprising the remaining genera of Nicotianeae plus the genus *Browallia* L. (tribe Salpiglossideae in D’Arcy, 1991; tribe Francisceae in Hunziker, 2001). This clade was provisionally called Petunioideae (Olmstead & al., 1999; validated by Thorne & Reveal, 2007), and subsequently called Petunieae (Olmstead & Bohs, 2007), which is the name used in most subsequent studies for this clade (e.g., Olmstead & al., 2008; Särkinen & al., 2013; Dupin & al., 2017; Huang & al., 2023; Pezzi & al., 2024). Plowman (1989) provided a taxonomic revision of the South American species of *Brunfelsia*.

> ***/Petuniina*** S.D. Smith & R.G. Olmstead, new clade name. **Regnum number**: 1111.— **Definition:** The minimum crown clade containing *Petunia axillaris* (Lamarck) Britton, Sterns, and Poggenburg 1888, *Fabiana imbricata* Ruiz & Pavón 1799, and *Calibrachoa parviflora* (Jussieu) D’Arcy 1989.

**Primary reference phylogeny:** This paper (Fig. 1A). See also this paper (Figs. 2, S1-S2); Olmstead & al. (2008: Fig. 1); Alaria & al. (2022: Fig. 1); Pezzi & al. (2024: Fig. 2). **Etymology**: From the genus *Petunia* named from the archaic French “petun” or Portuguese “petum”, believed derived from the indigenous Guarani for tobacco. **Composition:** *Petunia, Calibrachoa* Cerv., and *Fabiana* Ruiz & Pav. **Synonyms:** None. **Comments:** *Calibrachoa* has been included in *Petunia* (e.g., Hunziker, 2001), with *Fabiana* placed in the same tribe, along with *Nicotiana* and others. Molecular phylogenetic studies have shown the close relationship of these three genera to the exclusion of others and have suggested all of the three possible relationships among the three genera (Olmstead & al., 2008; Sarkinen & al., 2013; Mader & Freitas, 2019). However, recent studies have reached a consensus that *Fabiana* is more closely related to *Calibrachoa* than either is to *Petunia* (Alaria & al., 2022; Pezzi & al., 2024). Also supporting the separation of *Calibrachoa* from *Petunia* and its close relationship with *Fabiana* is the chromosome count of X=9 for *Calibrachoa* and *Fabiana* and X=7 for *Petunia* (Stehmann & al., 2009).

> ***/Schwenckieae*** Hunziker 1977 [R.G. Olmstead & R. Deanna] converted clade name. **Regnum number:** 1112.—**Definition:** The minimum crown clade containing *Schwenckia americana* Linnaeus 1764, *Heteranthia decipiens* Nees & Martius 1823, and *Melananthus guatemalensis* Solereder 1892.

**Primary reference phylogeny:** This paper (Fig. 1A). See also this paper (Figs. 2, S1-S2); Olmstead & al. (2008: Fig. 1). **Etymology:** From the genus *Schwenckia* L. named to honor Carl August von Schwenck, a German botanist. **Composition:** The genera *Heteranthia* Nees & Mart., *Melananthus* Walp., and *Schwenckia* L. **Diagnostic apomorphy:** Petal appendages composed of a median distal projection of the three-lobed petal occur in the genera *Melananthus* and *Schwenckia*, but are absent in *Heteranthia* (Paucar & al., 2020) apparently through loss. **Synonyms:** None. **Comments:** In traditional classifications (e.g., D’Arcy, 1991; Hunziker, 2001; Barboza & al., 2016), tribe Schwenckieae always includes *Schwenckia* and *Melananthus*, and occasionally includes *Heteranthia* and/or *Protoschwenkia*. *Protoschwenkia* was found to belong in Cestroideae by Olmstead & al. (2008).

> ***/Dodecachroma*** R.G. Olmstead & R. Deanna, new clade name. **Regnum number**: 1113.— **Definition:** The minimum crown clade containing *Solanum nigrum* Linnaeus 1753 and *Nicotiana tabacum* Linnaeus 1753.

**Primary reference phylogeny:** This paper (Fig. 1A). See also this paper (Figs. 2, S1-S2); Olmstead & al. (2008: Fig. 1). **Etymology**: From the putative synapomorphy for this clade of a base chromosome number of 12. **Composition:** /*Nicotianoideae* and /*Solanoideae*. **Diagnostic apomorphy**: A base chromosome number of X=12 (Olmstead & Palmer, 1992). **Synonyms:** The informal name “X=12 clade” has been used to refer to this clade (Olmstead & Sweere, 1994). **Comments:** Most traditional classifications of Solanaceae recognized two subfamilies, Cestroideae and Solanoideae. Solanoideae were recognized to have a common base chromosome number of X=12, but Cestroideae exhibited a wide range of base chromosome numbers from X=7 to X=12. Molecular phylogenetic studies discovered that Cestroideae, as traditionally circumscribed, were paraphyletic, with the X=12 representatives, *Nicotiana* L. and the Australian endemic tribe Anthocercideae, strongly supported as sister to Solanoideae (Olmstead & Sweere, 1994; Martins & Barkman, 2005; Olmstead & al., 2008; Ng & Smith, 2016). No morphological trait, other than chromosome number, has been identified that characterizes this clade.

> ***/Nicotianoideae*** Miers 1848 [R.G. Olmstead & R. Deanna] converted clade name. **Regnum number**: 1114.—**Definition:** The maximum crown clade containing *Nicotiana tabacum* Linnaeus 1753, but not *Solanum nigrum* Linnaeus 1753.

**Primary reference phylogeny:** This paper (Fig. 1A). See also this paper (Figs. 2, S1-S2); Olmstead & al. (2008: Fig. 1). **Etymology**: From the genus *Nicotiana* L. named for Jean Nicot, a 16th century diplomat who is said to have introduced the plant to France. **Composition:** Clade */Anthocercideae* and the genus *Nicotiana.* **Diagnostic apomorphy**: Members of /*Nicotianoideae* share a ca. 100 bp deletion in the *trnA* intron in the plastid genome (Olmstead & Palmer, 1992). **Synonyms:** None. **Comments:** Based primarily on fruit morphology, *Nicotiana* and the Australian tribe Anthocercideae were traditionally assigned to subfamily Cestroideae (e.g., D’Arcy, 1991). They share the base chromosome number X=12 and molecular phylogenetic studies have found them to belong together in a clade sister to subfamily Solanoideae (also X=12). However, no morphological synapomorphy is known for the clade. Nicotianoideae was first used for this clade by Olmstead & al. (1999). Names at the rank of family, subfamily, tribe, and subtribe all have been based on the genus *Nicotiana*, but none of them match the circumscription adopted here.

> ***/Anthocercideae*** G. Don 1838 [R.G. Olmstead & R. Deanna] converted clade name. **Regnum number**: 1115.—**Definition:** The maximum crown clade containing *Anthocercis littorea* Labillardière 1806, but not *Nicotiana tabacum* Linnaeus 1753.

**Primary reference phylogeny:** This paper (Fig. 1A). See also this paper (Figs. 2, S1-S2); Olmstead & al. (2008: Fig. 1). **Etymology**: From the genus *Anthocercis* Labill., from the Greek “anthoos” (flower) and “kerkis” for ray (narrow) in reference to the narrow corolla lobes. **Composition:** Tribe Anthocercideae, including *Anthocercis, Anthotroche* Endl.*, Crenidium* Haegi*, Cyphanthera* Miers*, Duboisia* R.Br.*, Grammosolen* Haegi, and probably *Symonanthus* Haegi. **Diagnostic apomorphy**: Corolla with “anthocercidoid” aestivation manifesting as each lobe with overlapping borders, two lobes right-involute, the remaining three left-involute (Hunziker, 2001). **Synonyms:** Subfamily *Anthocercidoideae* (G.Don) Tétényi. **Comments:** While this study found a monophyletic /*Anthocercideae*, including *Symonanthus* (Fig. 1, Supp. Fig. 1), the clade is either weakly supported (Clarkson & al., 2004), or *Symonanthus* is unresolved relative to *Nicotiana* and other /*Anthocercideae* in most prior studies (Garcia & Olmstead, 2003; Olmstead & al., 2008; Särkinen & al., 2013; Ng & Smith, 2016). There is good reason, however, to believe that *Symonanthus* belongs to this clade based on a combination of anatomical, pollen, and biochemical data (reviewed in Garcia & Olmstead, 2003), as well as biogeography, which indicates that *Nicotiana* originated in South America long after the origin of Anthocercideae, and only one highly derived clade within *Nicotiana* occurs in Australia (Clarkson & al., 2004; Olmstead, 2013; Dupin & al., 2017). If further work discovers that *Symonanthus* is not part of this clade, the clade definition adopted here will exclude it.

> ***/Solanoideae*** Burnett 1835 [R.G. Olmstead & R. Deanna] converted clade name. **Regnum number**: 1116.—**Definition:** The maximum crown clade containing *Solanum nigrum* Linnaeus 1753, but not *Nicotiana tabacum* Linnaeus 1753.

**Primary reference phylogeny:** This paper (Fig. 1A). See also this paper (Figs. 2, S1-S2); Olmstead & Palmer (1992: Fig. 3); Olmstead & al. (2008: Fig. 1). **Etymology**: From the genus *Solanum*. The name was first used by Pliny the Elder (Quattrocchi, 2000). The Latin derivation of the name may refer to “sol” (sun) in reference to the flower shape, or more likely to“solamen” or “solare “ (comfort, solace, or soothe) for the medicinal qualities of the plants. **Composition:** Clades /*Atropina, /Capsiceae, /Datureae, /Physalideae, /Solandreae, /Solaneae*, and the genera *Exodeconus* Raf.*, Mandragora* L.*, Nicandra* Adans., and *Salpichroa*. **Diagnostic apomorphy**: Fruit a berry, usually fleshy but occasionally modified (e.g., *Datura*, fleshy fruits also found in *Cestrum* but are modified capsules, Pabón-Mora & Litt, 2011) and discoidal-reniform, campylotropous, flattened seeds with curved embryos (Särkinen & al., 2018a). **Synonyms:** None. **Comments:** */Solanoideae* has long been recognized as a cohesive group within Solanaceae with a base chromosome number of X=12 and consistent floral, seed, and fruit traits.

> ***/Atropina*** Olmstead & al. 2008 [R.G. Olmstead & R. Deanna] converted clade name. **Regnum number**: 1118.—**Definition:** The minimum crown clade containing *Atropa bella-donna* Linnaeus 1753, *Hyoscyamus niger* Linnaeus 1753, *Lycium afrum* Linnaeus 1753, *Jaborosa integrifolia* Lamarck 1789, *Latua pubiflora* (Grisebach) Baillon 1888.

**Primary reference phylogeny:** This paper (Fig. 1A). See also this paper (Figs. 2, S1-S2); Olmstead & al. (2008: Fig. 1); Chiarini & al. (2022: Fig. 2). **Etymology**: From the genus *Atropa* L. named for one of the three Greek fates, Atropos, who snipped the thread of life of mortals, perhaps for the well-known lethal effects of the alkaloid components of the plant *Atropa bella-donna* L. **Composition:** The clades */Hyoscyameae, /Lyciina,* and genera *Jaborosa* Juss. and *Latua* Phil. **Synonyms:** None. **Comments:** Atropina was first coined as an informal clade name by Olmstead & al. (2008) and used in subsequent phylogenetic studies of Solanaceae (Särkinen & al., 2013; Dupin & al., 2017). A close relationship among members of this clade was not predicted by traditional classifications, which often excluded *Nolana* L.f. and *Sclerophylax* Miers from the family Solanaceae.

> ***/Hyoscyameae*** Endlicher 1839 [R.G. Olmstead & T. Särkinen] converted clade name. **Regnum number**: 1119.—**Definition:** The minimum crown clade containing *Atropa bella-donna* Linnaeus 1753 and *Hyoscyamus niger* Linnaeus 1753.

**Primary reference phylogeny:** This paper (Fig. 1A). See also this paper (Figs. 2, S1-S2); Yuan & al. (2006: Fig. 2); Olmstead & al. (2008: Fig. 1); Tu & al. (2020: Fig. 1); Lei & al. (2021: Fig. 2). **Etymology**: From the genus *Hyoscyamus* L.; the name was given by the 1st Century Greek physician Dioscorides from the Greek “hyoskyamos” (hog-bean). **Composition:** /*Hyoscyaminae* and the genus *Atropa.* **Synonyms:** None. **Comments:** Tribe Hyoscyameae and subtribe Hyoscyaminae have both been used for a taxon defined here as /*Hyoscyaminae*. *Atropa* is sister to the rest of this clade and differs primarily in its fruit, having a fleshy berry similar to other Solanoideae, whereas the others have a distinctive circumscissile capsule-like berry. Primarily for this reason, *Atropa* has been assigned elsewhere in classifications prior to molecular phylogenetic studies (e.g, D’Arcy, 1991; Hunziker, 2001). However, early molecular phylogenetic studies (Olmstead & Palmer, 1992; Olmstead & al., 1999; Yuan & al., 2006; Olmstead & al., 2008) identified a strongly supported clade that included the genus *Atropa* with the traditionally circumscribed tribe Hyoscyameae, but without strong internal support and expanded the concept of the tribe to include *Atropa* (see also Barboza & al., 2016). We adopt this circumscription for /*Hyoscyameae*, and use the narrower concept for /*Hyoscyaminae* that excludes *Atropa* (see below).

> **/*Hyoscyaminae*** Kittel 1840 [T. Särkinen & R.G. Olmstead] converted clade name. **Regnum number**: 1120.—**Definition:** The maximum crown clade containing *Hyoscyamus niger* Linnaeus 1753, but not *Atropa bella-donna* Linnaeus 1753.

**Primary reference phylogeny:** This paper (Fig. 1A). See also this paper (Figs. 2, S1-S2); Olmstead & al. (2008: Fig. 1); Yuan & al. (2006: Fig. 2); Hajrasouliha & al. (2014: Fig. 3); Sanchez-Puerta & Abbona (2014: Fig. 3); Tu & al. (2020: Fig. 1); Lei & al. (2021: Fig. 2). **Etymology**: From the genus *Hyoscyamus* L.; the name was given by the 1st Century Greek physician Dioscorides from the Greek “hyoskyamos” (hog-bean). **Composition:** The genera *Atropanthe* Pascher*, Anisodus* Link & Otto*, Hyoscyamus* L.*, Physochlaina* G.Don*, Przewalskia* Maxim.*, Scopolia* Jacq. **Diagnostic apomorphy**: Fruit a circumscissile capsule-like berry (pyxidium). **Synonyms:** Tribe Hyoscyameae sensu Hunziker (2001). **Comments:** This is a cohesive clade long recognized in traditional classifications due to the distinctive circumscissile dehiscence of the dry berry, often referred to as a capsule. However, differences in rank to which the group is assigned (e.g., tribe and subtribe) and circumscription of the included genera have varied among treatments (Lu & Zhang, 1986; Hoare & Knapp, 1997; Hunziker, 2001). Phylogenetic studies of /*Hyoscyaminae* indicate strong support for the clade, but also conflict regarding generic circumscriptions and relationships (see secondary reference trees cited above). For that reason, we provide a maximum clade definition.

> ***/Lyciina*** Olmstead & Bohs, 2007 [R.G. Olmstead & R. Deanna] converted clade name. **Regnum number**: 1121.—**Definition:** The minimum crown clade containing *Lycium afrum* Linnaeus 1753, *Nolana humifusa* (Gouan) I. M. Johnston 1936, and *Sclerophylax spinescens* Miers 1848.

**Primary reference phylogeny:** This paper (Fig. 1A). See also this paper (Figs. 2, S1-S2); Olmstead & al. (2008: Fig. 1); Chiarini & al. (2022: Fig. 2). **Etymology**: From the genus *Lycium* L. named following the use of this name by Dioscorides for a plant from the region of Greece known as Lycia. **Composition:** The genera *Lycium, Nolana* L. and *Sclerophylax* Griseb. **Synonyms:** None. **Comments:** Lycieae has been used as a tribe in recent classifications (D’Arcy, 1991; Hunziker, 2001; Barboza & al., 2016) and phylogenetic studies (e.g., Olmstead & al., 2008; Levin & al., 2009, 2011) to refer to a group of three genera *Lycium*, *Grabowskia* Schltdl., and *Phrodus* Miers. These phylogenetic studies have shown that *Grabowskia* is nested within *Lycium* and *Phrodus* has a conflicting placement in plastid vs. nuclear gene trees, so all three genera are now included in *Lycium* (Barboza & al., 2016). *Nolana* and *Sclerophylax* were either not included in Solanaceae in traditional treatments (e.g., Hunziker, 2001) or, if included, not associated with Lycieae (e.g., D’Arcy, 1991; Barboza & al., 2016), but have been found to be well-supported as part of a clade with tribe Lycieae. The inclusive clade was given the informal name Lyciina by Olmstead & Bohs (2007). Lyciinae, Lycieae, and Lycioideae are all available to be converted clade names, but all differ significantly by exclusion of *Nolana* and *Sclerophylax*, so Lyciina is adopted here to avoid confusion with the traditional use of Lycieae.

> ***/Solandreae*** Miers 1849 [A. Orejuela & R.G. Olmstead] converted clade name. **Regnum number**: 1122.—**Definition:** The minimum crown clade containing *Solandra grandiflora* O. Swartz 1787, *Schultesianthus leucanthus* (J. Donnell Smith) A.T. Hunziker 1977, *Juanulloa parasitica* Ruiz & Pavón 1794, and *Doselia lopezii* (A.T. Hunziker) A. Orejuela & Särkinen 2022.

**Primary reference phylogeny:** This paper (Fig. 1A). See also this paper (Figs. 2, S1-S2); Olmstead & al. (2008: Fig. 1); Orejuela & al. (2017: Fig. 2). **Etymology**: From the genus *Solandra* Sw. named for Daniel Carl Solander, 18th century Swedish botanist and pupil of Linnaeus. **Composition:** The genera *Doselia* Orejuela & Särkinen, *Dyssochroma* Miers*, Ectozoma* Miers*, Hawkesiophyton* Hunz.*, Juanulloa* Ruiz & Pav.*, Markea* A.Rich.*, Merinthopodium* Donn.Sm.*, Poortmannia* Drake*, Rahowardiana* D’Arcy*, Schultesianthus* Hunz.*, Solandra,* and *Trianaea* Planch. & Linden. **Diagnostic apomorphy**: Epiphytes or woody climbers with tubular flowers and fleshy to leathery berries. **Synonyms:** None. **Comments:** Solandreae and Juanulloeae often have been considered separate tribes (D’Arcy, 1991; Hunziker, 2001; Barboza & al., 2016). Knapp & al. (1997) found *Solandra* nested within a Juanulloeae in a morphology-based phylogeny. Molecular phylogenies consistently obtained this clade, albeit with weak support, starting with Olmstead & Sweere (1994), sometimes with *Solandra* nested within Juanulloeae (e.g., Olmstead & al., 2008) and sometimes as sister to it (e.g., Orejuela & al., 2017; Huang & al. 2023). The present definition follows a clade consistently retrieved in recent molecular phylogenies and recognises the monophyly of the group including both *Solandra* and the historically defined Juanulloeae.

> ***/Solaneae*** Dumortier 1829 [R.G. Olmstead & R. Deanna] converted clade name. **Regnum number**: 1117.—**Definition:** The minimum crown clade containing *Datura stramonium* Linnaeus 1753, *Physalis pubescens* Linnaeus 1753, *Solanum nigrum* Linnaeus 1753, *Jaltomata procumbens* (Cavanilles) J.L. Gentry 1973, *Salpichroa glandulosa* (W.J. Hooker) Miers 1845, and *Capsicum annuum* Linnaeus 1753.

**Primary reference phylogeny:** This paper (Fig. 1A). See also this paper (Figs. 2, S1-S2); Olmstead & Palmer (1992: Fig. 2); Olmstead & al. (2008: Fig. 1); Poczai & Hyvönen (2013: Fig. 1). **Etymology**: For the position of this clade within the larger Solanoideae as a monophyletic group. **Composition:** */Capsiceae, /Datureae, /Physalideae,* and the genera *Solanum* and *Jaltomata.* **Diagnostic apomorphy**: Presence of *trnF* duplicate pseudogene (Poczai & Hyvönen, 2013). **Synonyms:** Pseudosolanoideae (Poczai & Hyvönen, 2013). **Comments:** A taxon closely approximating the composition of this clade has been recognized at the rank of tribe in most traditional classifications of Solanaceae (e.g., D’Arcy, 1991; Hunziker, 2001). Those taxa often included the genera *Atropa*, *Exodeconus*, and *Mandragora* that are excluded from this clade, and did not include tribe Datureae that is included here. This clade was recognized as Solaneae in the earliest molecular phylogeny of Solanaceae by Olmstead & Palmer (1992), but in the interest of creating a series of monophyletic tribes within subfamily Solanoideae, Solaneae was subsequently restricted to a clade composed of *Solanum* and *Jaltomata* (Olmstead & al., 1999, 2008; Sarkinen & al., 2013; Barboza & al., 2016). All of the early studies were based on plastid DNA, but recent studies have documented discordance between plastid and nuclear trees (Wu & al., 2019; Powell & al., 2022; Huang & al., 2023) that calls into question whether *Solanum* and *Jaltomata* are a clade. So, we recognize a clade /*Solaneae* that corresponds closely to tribe Solaneae of traditional classifications and exactly to the first phylogenetic application of the name (Olmstead & Palmer, 1992).

> ***/Datureae*** Dumortier 1829 [S.D. Smith & R.G. Olmstead] converted clade name. **Regnum number**: 1123.—**Definition:** The maximum crown clade containing *Datura stramonium* Linnaeus 1753, but not *Solanum nigrum* Linnaeus 1753, *Capsicum annuum* Linnaeus 1753, *Jaltomata procumbens* (Cavanilles) J.L. Gentry 1973, *Salpichroa glandulosa* (W.J. Hooker) Miers 1845, and *Physalis angulata* Linnaeus 1753.

**Primary reference phylogeny:** This paper (Fig. 1A). See also this paper (Figs. 2, S1-S2); Olmstead & al. (2008: Fig. 1); Dupin & Smith (2018: Fig. 3). **Etymology**: From the genus *Datura* L. named from the Hindu “dhatura” meaning “thorn-apple.” **Composition:** The genera *Brugmansia* Pers., *Datura*, and *Trompettia* J.Dupin. **Diagnostic apomorphy**: Flowers with contorted-conduplicate corolla aestivation (Hunziker, 2001). **Synonyms:** None. **Comments:** *Brugmansia* has been included in *Datura* or more recently maintained as a separate genus (Persson & al., 1999; Hay & al., 2012), but their close relationship has not been in doubt. The type species of genus *Brugmansia* (*B.* ✕ *candida)* is of hybrid origin, with *B. aurea*, the species included in the reference phylogeny, one of its parents (Hay & al., 2012). *Trompettia cardenasiana*, originally named in distantly related *Iochroma* Benth., was found to be sister to *Datura* and *Brugmansia* (Olmstead & al., 2008) and included in tribe Datureae (Dupin & Smith, 2018).

> ***/Physalideae*** Reveal 2012 [R.G. Olmstead & R. Deanna] converted clade name. **Regnum number**: 1124.—**Definition:** The maximum crown clade containing *Physalis pubescens* Linnaeus 1753 *(Physalis angulata* Linnaeus 1753), but not *Salpichroa glandulosa* (W.J. Hooker) Miers 1845, *Capsicum annuum* Linnaeus 1753, *Jaltomata procumbens* (Cavanilles) J.L. Gentry 1973, *Datura stramonium* Linnaeus 1753, and *Solanum nigrum* Linnaeus 1753.

**Primary reference phylogeny:** This paper (Fig. 1). See also this paper (Figs. 2, S1); Olmstead & al. (2008: Fig. 1); Deanna & al. (2019: Fig. 2); Sandoval-Padilla & al. (2025: Fig. 3). **Etymology**: From the genus *Physalis* L. named from the Greek “phusallis” meaning bladder, a reference to the inflated calyx. **Composition:** Subtribes Physalidinae, Iochrominae, and Withaninae, and the genera *Cuatresia* Hunz. and *Deprea* Raf. (Olmstead & al., 2008; Deanna & al., 2019). **Synonyms:** Physaleae (see comments). “Physaloids” has been used as an informal name for this clade. **Comments:** Miers (1849) named the tribe Physaleae, which was corrected to Physalideae by Reveal (2012) to conform to the rules of the *Code* (Turland & al., 2018). Physaleae was first used for this clade by Olmstead & al. (1999) and was used by subsequent authors (Olmstead & al., 2008; Särkinen & al., 2013; Dupin & al., 2017), but subsequent to publication of the corrected spelling, Physalideae has been used by Deanna & al. (2019), Huang & al. (2023), and in the global treatment of Solanaceae by Barboza & al. (2016), so we use /*Physalideae* to provide continuity of use. The monophyly of /*Physalideae* is not in doubt, but internal relationships are uncertain and not consistent across published phylogenies (e.g, Zamora-Tavares & al., 2016; Deanna & al., 2019; Sandoval-Padilla & al., 2025), therefore, additional work is needed before clades within /*Physalideae* can be defined with confidence. The type species of the genus *Physalis*, *P. pubescens*, is not included in the primary reference tree, but is most closely related to *P. angulata* in the phylogeny of Deanna & al. (2019).

> ***/Capsiceae*** Dumortier 1827 [R.G. Olmstead & R. Deanna] converted clade name. **Regnum number**: 1125.—**Definition:** The minimum crown clade containing *Capsicum annuum* Linnaeus 1753, *Lycianthes lycioides* (Linnaeus) Hassler 1917, and *Lycianthes biflora* (Loureiro) Bitter 1919.

**Primary reference phylogeny:** This paper (Fig. 1A). See also this paper (Figs. 2, S1-S2); Olmstead & al. (2008: Fig. 1). **Etymology**: From the genus *Capsicum* L. named from the Greek “kapto” (to bite), or the Latin “capsa” (box); perhaps referring to the pungency or shape of the fruit. **Composition:** The genera *Capsicum* and *Lycianthes* (Dunal) Hassl. **Diagnostic apomorphy**: A calyx with ten primary vascular traces, as opposed to five in most Solanaceae; these traces often manifest as submarginal appendages. **Synonyms:** None. **Comments:** *Lycianthes* shares the distinctive trait of poricidal anthers with *Solanum* L., leading to the previous conclusion of a close relationship between the two, despite sharing the shared calyx trait with *Capsicum*. Phylogenetic studies provide strong support for /*Capsiceae* with *Capsicum* derived from a paraphyletic *Lycianthes* (Spalink & al., 2018). A complete taxonomic treatment of *Capsicum* has been provided by Barboza & al. (2022).

> ***/Solanum*** Linnaeus 1753 [L. Bohs, S. Knapp, L.L. Giacomin & T. Särkinen] converted clade name. **Regnum number**: 1126.—**Definition:** The maximum crown clade containing *Solanum nigrum* Linnaeus 1753, but not *Jaltomata procumbens* (Cavanilles) J.L. Gentry 1973, *Capsicum annuum* Linnaeus 1753, *Salpichroa origanifolia* (Lamarck) Baillon 1888, *Datura stramonium* Linnaeus 1753, and *Physalis pubescens* Linnaeus 1753.

**Primary reference phylogeny:** This paper (Fig. 1A). See also this paper (Figs. 2, S1, S2); Olmstead & al. (2008: Fig. 1); Gagnon & al. (2022: Fig. S-11); Messeder & al. (2024: Fig, 1). **Etymology**: From the genus *Solanum*. The name was first used by Pliny the Elder (Quattrocchi, 2000). The Latin derivation of the name may refer to “sol”(sun) in reference to the flower shape, or more likely to“solamen” or “solare “ (comfort, solace, or soothe, for the medicinal qualities of the plants). **Composition:** The clade represents the genus *Solanum*, which contains 1,239 species distributed worldwide (Hilgenhof & al., 2023) and represents approximately half the species diversity of the family. **Diagnostic apomorphy**: Recognized by the combined set of characters of anthers dehiscing by apical pores or pores lengthening to slits, usually 5-merous flowers with 5 vascular traces in the calyx, and berry fruits. **Synonyms:** None. **Comments:** *Solanum* is the most species-rich genus in the Solanaceae and contains many taxa previously recognized at the generic level (e.g., *Aquartia* Jacq., *Cyphomandra* Sendtn.*, Lycopersicon* Mill.*, Normania* Lowe and *Triguera* Cav.). An overview of the lineages recognized, as well as morphology and geography, can be found in Gagnon & al. (2002) and Hilgenhof & al. (2023).

> **/*Thelopodium*** Bohs 2005 [L. Bohs & S. Knapp] converted clade name. **Regnum number**: 1127.—**Definition:** The minimum crown clade containing *Solanum thelopodium* Sendtner 1846 and *Solanum monarchostemon* S. Knapp 2000.

**Primary reference phylogeny:** This paper (Fig. 1B). See also this paper (Fig. S1); Bohs (2005: Fig. 1). **Etymology**: From the epithet of *Solanum thelopodium* Sendtn., the first described member of the clade. **Composition:** Three species of slender, single-stemmed understory shrubs from tropical America. **Diagnostic apomorphy**: Tapered and strongly unequal anthers (one long, two medium, and two short). **Synonyms:** none. **Comments:** Bohs (2005) identified *S. theopodium* as the first branching lineage in *Solanum*, a position confirmed by all subsequent phylogenies that have included other species (e.g., Särkinen & al. 2013; Gagnon & al. 2022; Messeder & al. 2024). A taxonomic revision of this clade (as the “*Solanum thelopodium* species group”) was published by Knapp (2000).

> **/*Regmandra*** Dunal 1852 [L. Bohs & T. Särkinen] converted clade name. **Regnum number**: 1128.—**Definition:** The minimum crown clade containing *Solanum multifidum* Lamarck 1794 and *Solanum montanum* Linnaeus 1753.

**Primary reference phylogeny:** This paper (Fig. 1B). See also this paper (Fig. S1); Bohs (2005: Fig. 1). **Etymology**: From a named (grad. ambig. “*Regmandra*”), but unranked, group of species in Dunal (1852). **Composition:** Twelve species of herbs from lomas vegetation (i.e., coastal fog forests) in the western Andean slopes in Peru and Chile. **Diagnostic apomorphy**: Most species with expanded stigmas and fruits with stone cells, neither of which are restricted to this clade. **Synonyms:** *Solanum* section *Regmandra* (Dunal) Ugent. **Comments:** The taxonomy of **/***Regmandra* was revised by Bennett (2008). Tepe & al. (2016) included **/***Regmandra* within **/***Pinnatifida* (see below) but Gagnon & al. (2022) showed nuclear-plastome discordance in the placement of the group within *Solanum* and recognized **/***Regmandra* as a distinct major clade in the genus.

> /***Peregrina*** S. Knapp & E. Gagnon, new clade name. **Regnum number**: 1129.—**Definition:** The maximum crown clade containing *Solanum terminale* Forsskål 1775, but not *Solanum montanum* Linnaeus 1753, *Solanum nigrum* Linnaeus 1753, *Solanum dulcamara* Linnaeus 1753, and *Solanum tuberosum* Linnaeus 1753.

**Primary reference phylogeny:** This paper (Fig. 1B). See also this paper (Fig. S1); Gagnon & al. (2022: Fig. S12); Messeder & al. (2024: Fig. 1a). **Etymology**: Derived from the Latin “peregrinus” for strange, in reference to the unusual geographical distribution and morphological variability of groups within this clade. **Composition:** Approximately 30 species of herbs, shrubs and woody vines with widely disparate distributions; species occur in southern South America, Australia (incl. New Guinea), continental Africa, Madagascar, and Macaronesia. **Diagnostic apomorphy**: None. **Synonyms:** “VANans” is an informal name applied to this clade by Gagnon & al. (2022). **Comments:** This clade was first recognised by Gagnon & al. (2022) and there was given the informal name of “VANAns”-an acronym composed from letters representing the main taxa involved. /*Peregrina* is a heterogenous group composed of several previously recognised groups, plus two species, *Solanum valdiviense* and the related *S. alphonsei* DC. (not included in the primary reference phylogeny but treated in Gagnon & al., 2022), that had previously been treated as members of /*Dulcamara* (as the Dulcamaroid clade) by Knapp (2013). Both are semi-woody vines or lax shrubs from the southern temperate forests of Chile and adjacent Argentina. Previously recognised groups included in /*Peregrina* are *S*. section *Archaeosolanum* (Marzell) Danert, comprising eight species of soft shrubs endemic to Australasia (incl. New Guinea and New Zealand) commonly known as the kangaroo apples, members of the group of shrubs and large woody vines from Madagascar and continental Africa called the “African non-spiny” clade by Knapp (2016) and the two species from *Solanum* section *Normania* (Lowe) Bitter (*S. trisectum* Dunal, *S. nava* Webb. & Berthel.) plus the previously recognised genus *Triguera* Cav. (*S. herculeum* Bohs) that were shown to be nested in *Solanum* by Bohs & Olmstead (2001). The monophyly of /*Peregrina* is supported by various phylogenomic analyses of different gene regions, both nuclear and plastid (e.g., Gagnon & al., 2022; Messeder & al., 2024; this paper). Taxonomic treatment of the African and Madagascan species was provided by Knapp (2016) and the kangaroo apples were treated taxonomically by Symon (1994) and phylogenetically by Poczai & al. (2011). The morphology, taxonomy, and relationships of the Macaronesian species (i.e., section *Normania*) were examined in detail in Francisco-Ortega & al. (1993) and these plus *S. herculeum* in Bohs & Olmstead (2001). Descriptions of the morphology of these groups can also be found in Hilgenhof & al. (2023).

> ***/Pinnatifida*** S. Knapp & L. Bohs, new clade name. **Regnum number**: 1130.—**Definition:** The minimum crown clade containing *Solanum tuberosum* Linnaeus 1753, *Solanum lycopersicum* Linnaeus 1753, *Solanum muricatum* Aiton 1789, *Solanum etuberosum* Lindley 1834, *Solanum sodiroi* Bitter 1912, *Solanum suaveolens* Kunth & C.D. Bouché 1848, *Solanum sanctae-marthae* Bitter 1913, *Solanum oxycoccoides* Bitter 1919, *Solanum mite* Ruiz & Pavón 1799, and *Solanum trifolium* Dunal 1852.

**Primary reference phylogeny:** This paper (Fig. 1B). See also this paper (Fig. S1); Tepe & al. (2016: Fig. 2). **Etymology**: From the common vegetative characteristics of species in this clade, most of which have variously pinnate to pinnatifid leaves. **Composition:** Approximately 186 non-prickly herbs to weakly woody species all from the Americas, with several widely cultivated (e.g., *S. tuberosum* L., *S. lycopersicum* L., *S. muricatum* Aiton). **Diagnostic apomorphies**: Most species have pinnatifid leaves and well-developed pseudostipules. **Synonyms:** Informally called “Potato” or “Potato clade” in previous publications (i.e., Tepe & al., 2016; Gagnon & al. 2022). **Comments:** This clade comprises all the wild relatives of the cultivated potato (*S. tuberosum*), tomato (*S. lycopersicum*), and pepino (S*. muricatum*) plus a number of other groups previously thought related (e.g., D’Arcy, 1972) such as *Solanum* section *Pteroidea* Dunal and *Solanum* section *Herpystichum* Bitter. Tepe & al. (2016) include **/***Regmandra* within **/***Pinnatifida* but Gagnon & al. (2022) showed nuclear-plastome discordance in the placement of **/***Regmandra* within *Solanum*.

> **/*Lycopersicon*** Miller 1768 [R.G. Olmstead & L. Bohs] converted clade name. **Regnum number**: 1131.—**Definition:** The minimum crown clade containing *Solanum lycopersicum* Linnaeus 1753 and *Solanum lycopersicoides* Dunal 1852.

**Primary reference phylogeny:** This paper (Fig. 1B). See also this paper (Fig. S1); Gagnon & al. (2022: Fig. S12). **Etymology**: From the generic name coined by Philip Miller in 1768 to segregate a group of taxa including *Solanum lycopersicum* L. as a distinct genus, referring to the earlier pre-Linnaean Tournefortian name for the tomato meaning “wolf peach” (see Peralta & al., 2008). **Composition:** Seventeen species of herbs, herbaceous vines and small shrubs all from Andean South America; one species (*S*. *lycopersicum*) is widely cultivated and often escapes and naturalizes. **Diagnostic apomorphies**: All species with yellow flowers and pinnatifid leaves, most (see comments) with elongate terminal anther appendices. **Synonyms:** *Solanum* section *Lycopersicon* (Mill.) Wettst.; *Solanum* subgenus *Lycopersicon* (Mill.) Seithe. **Comments:** /*Lycopersicon* contains all crop wild relatives of the cultivated tomato, *S. lycopersicum*. Species treated as members of section *Lycopersicon* by Peralta & al. (2008) have beak-like, elongate terminal anther appendages, while the rest of the species in /*Lycopersicon* have ellipsoid anthers without elongate beaks. These have been treated as *Solanum* sections *Lycopersicoides* (A.Child) Peralta and *Juglandifolia* (Rydb.) A.Child (Peralta & al., 2008). Because *S. lycopersicum* is a major crop, a wealth of genomic resources are available for both the clade (e.g., Gao & al., 2019; Zhou & al., 2022; Li & al., 2023).

> **/*Petota*** Dumortier 1827 [L. Bohs & R.G. Olmstead] converted clade name. **Regnum number**: 1132.—**Definition:** The minimum crown clade containing *Solanum tuberosum* Linnaeus 1753 and *Solanum piurae* Bitter 1916.

**Primary reference phylogeny:** This paper (Fig. 1B). See also this paper (Fig. S1); Yan & al. (2022: Fig. 3A); Bozan & al. (2023: Fig. 3) **Etymology**: The origin of the name is not known, but is perhaps a Latinization of a common name for *S. tuberosum* L., the cultivated potato. **Composition:** A clade of 112 species all native to the Americas; four species are cultivated (*S. tuberosum* worldwide; *S. ajanhuiri* Juz., *S. juzepzcukii* Bukasov and *S. curtilobum* Juz. & Bukasov only in the high Andes of Peru and Bolivia). **Diagnostic apomorphies**: Herbaceous plants with pinnatifid leaves and tubers borne on underground stems. **Synonyms:** None. **Comments:** D’Arcy (1972) lists 35 different sectional and series names that he considered to belong to *Solanum* section *Petota* Dumort. Phylogenetic studies have shown these to be non-monophyletic (see Spooner & al., 2014). There are two parallel and complementary taxonomic systems used for */Petota*. The traditional Ochoa and Hawkes systems classify the clade based on morphology and cytogenetic data with the aim to recognise species based on distinct and useful tuber traits (232 species; Hawkes, 1990; Ochoa 1990, 1999). Spooner’s more recent phylogenetically-based system is based on a combination of morphological, cytogenetic, and molecular data with the aim to recognise evolutionarily distinct species (112 species; Spooner & al., 1993, 2004, 2007, 2014, 2016, 2019; Huamán & Spooner, 2002; Ovchinnikova & al., 2011). Because *S. tuberosum* is the world’s most important non-grain crop, a wealth of germplasm (e.g., Sotomayor & al., 2023) and genomic resources are becoming available for both the cultivated and wild species to assist in crop improvement (e.g., Bozan & al., 2023; Wu & al., 2023; Sun & al., 2025; Zhu & al., 2025).

> **/*Dulcamara*** Moench 1794 [S. Knapp & L. Bohs] converted clade name. **Regnum number**: 1133.—**Definition:** The maximum crown clade containing *Solanum dulcamara* Linnaeus 1753, but not *Solanum nigrum* Linnaeus 1753.

**Primary reference phylogeny:** This paper (Fig. 1B). See also this paper (Fig. S1); Bohs (2005: Fig. 1). **Etymology**: From the Latin “dulcis” (sweet) and “amar” (bitter), hence the English common name bittersweet, in reference to medicinal use of the bark (Dunal, 1813). This was first used by Linnaeus for *Solanum dulcamara* L., one of the most widespread species of the clade, and subsequently for the genus *Dulcamara* Moench. **Composition:** About 44 species of mostly vining or scandent shrubs distributed in the Americas, Asia, and Europe. **Diagnostic apomorphies**: All species of the clade have a cup-shaped pedicel base. **Synonyms:** *Solanum* section *Dulcamara* Dumort., *Solanum* subsection *Dulcamara* (Dumort.) Dunal. **Comments:** Moench (1794) coined the genus name *Dulcamara* to distinguish what he considered the very distinct *S. dulcamara*, based on its fused anthers. Traditional circumscriptions of this group have only included those species closely related to the European *S. dulcamara*. Other species included in /*Dulcamara* have been treated (D’Arcy, 1972) as *Solanum* section *Jasminosolanum* Seithe or *Solanum* section *Holophylla* (G.Don) Walp. (see Knapp, 2013 for history). Knapp (2013) provided a taxonomic revision of this clade including *S. valdiviense* Dunal, *S. alphonsei* Dunal and *S. salicifolium* Phil. as members. These three species are excluded from /*Dulcamara* here. *Solanum salicifolium* is now recognised as a member of /*Morella* (Gagnon & al., 2022; Knapp & al., 2023), and *S. alphonsei* and *S. valdiviense* form a clade informally named as Valdiviense. *Solanum valdiviense* forms part of /*Peregrina* here. Relationships amongst these taxa at the clade level are still unclear so we refrain from naming these informally recognised groups here.

> **/*Morella*** Dumortier 1827 [T. Särkinen & S. Knapp] converted clade name. **Regnum number**: 1134.—**Definition:** The maximum crown clade containing *Solanum nigrum* Linnaeus 1753, but not *Solanum dulcamara* Linnaeus 1753.

**Primary reference phylogeny:** This paper (Fig. 1B). See also this paper (Fig. S1); Bohs (2005: Fig. 1); Särkinen & al. (2015: Fig. 2); Gagnon & al. (2022: Fig. S12). **Etymology**: From “morelle” the common name of the black nightshade (*S. nigrum* L.) in Francophone Europe. **Composition:** A group of 78 non-prickly, herbaceous to weakly woody species distributed globally, with centres of diversity in the tropical Andes and Africa. **Diagnostic apomorphies**: Lacking the cup-shaped pedicel bases of /*Dulcamara*, except for *S. salicifolium* Phil. Some species with stone cells in fruits. **Synonyms:** “Morelloid” is an informal name used for this clade (Bohs, 2005; Sarkinen & al., 2015; Gagnon & al., 2022). **Comments:** *Solanum* section *Morella* Dumort. included the type species of the genus, so under the rules of traditional nomenclature was corrected to *Solanum* section *Solanum*. The clade comprises the traditional *Solanum* section *Solanum* with the addition of species traditionally (D’Arcy, 1972) recognized as *Solanum* sections *Campanulisolanum* Bitter *Chamaesarachidium* Bitter, and *Episarcophyllum* Bitter. Särkinen & al. (2018b), and Knapp & al. (2019, 2023) provided taxonomic revisions for the clade for the Eastern Hemisphere, North and Central America (incl. Caribbean), and South America, respectively. *Solanum salicifolium* is included in /*Morella* as defined here (see also under /*Dulcamara*).

> **/*Brevantherum*** Seithe 1962 [L.L. Giacomin & L. Bohs] converted clade name. **Regnum number**: 1135.—**Definition:** The maximum crown clade containing *Solanum erianthum* D. Don 1825 but not *Solanum betaceum* Cavanilles 1799, *Solanum graveolens* Bunbury 1841, *Solanum aphyodendron* S. Knapp 1985, *Solanum havanense* Jacquin 1794, and *Solanum wendlandii* J.D. Hooker 1887.

**Primary reference phylogeny:** This paper (Fig. 1B). See also this paper (Fig. S1); Bohs (2005: Fig. 1); Gagnon & al. (2022: Fig. S12); Messeder & al. (2024: Fig. 1b). **Etymology**: From the sectional name *Brevantherum* Seithe, which is a reference to the short, ellipsoid-oblong anthers. **Composition:** The clade includes ca. 90 species of herbs, shrubs, and trees distributed in the Americas, with three species becoming invasive in Africa, Asia, and Australia. **Diagnostic apomorphies**: Members have the unique character combination of a lack of prickles, stellate or lepidote trichomes (most species), and oblong anthers. **Synonyms:** *Solanum* subgenus *Brevantherum* (Seithe) D’Arcy. **Comments:** The limits of this clade have been explored and expanded from the narrower definition of *Solanum* section *Brevantherum* Seithe as a result of recent phylogenetic studies (e.g., Tovar & al., 2021; Gagnon & al., 2022; Messeder & al., 2024). Detailed taxonomic revisions and phylogenies have been published for some minor clades within /*Brevantherum*: e.g., section *Gonatotrichum* Bitter by Stern & Bohs (2012) and Stern & al. (2013); Erianthum and Abutiloides clades by Tovar & al. (2021) and Roe (1972), respectively. An expanded sampling using transcriptomics has been presented by Messeder & al. (2024). What was recognised in the past as *Solanum* sect. *Brevantherum* Seithe does not accommodate the diversity currently understood in /*Brevantherum*, which is closer to what was treated as *Solanum* subgenus *Brevantherum* (Seithe) D’Arcy (D’Arcy, 1972), with the addition of at least two clades that are mainly of herbs to small shrubs with unbranched trichomes. The current circumscription includes species related to *Solanum bradei* Giacomin & Stehmann and *Solanum* sect. *Gonatotrichum* (sensu Stern & al., 2013). A complete list of species is available in Hilgenhof & al. (2023).

> **/*Cyphomandra*** Sendtner 1845 [L. Bohs] converted clade name. **Regnum number**: 1136.— **Definition:** The maximum crown clade containing *Solanum betaceum* Cavanilles 1799, but not *Solanum aphyodendron* S.Knapp 1985, *Solanum erianthum* D.Don 1825, and *Solanum wendlandii* J.D. Hooker 1887.

**Primary reference phylogeny:** This paper (Fig. 1B). See also this paper (Fig. S1); Bohs (2005: Fig. 1). **Etymology**: From the genus *Cyphomandra* derived from the Greek “kyphos” (bent or humped) and “andros” (man), referring to the thickened anther connectives in species of the genus. **Composition:** Approximately 50 species of shrubs and small trees native to the American tropics. *Solanum betaceum* Cav. is the source of the cultivated tree tomato, also known as tomate de árbol or tamarillo. **Diagnostic apomorphies**: Very large chromosomes and nuclear DNA amounts [Chiarini & al. (2018), Deanna & al. (2022), Mesquita & al. (2025)]. Enlarged anther connectives in members of the former genus *Cyphomandra*. **Synonyms:** None. **Comments:** D’Arcy (1972) treated some of the species included here as the distinct genus *Cyphomandra* Sendtn. He treated other species included here as *Solanum* section *Cyphomandropsis* Bitter. *Solanum graveolens* Bunbury groups with */Cyphomandra* in our analysis here, but this placement varies or is poorly supported in others (e.g., Gagnon & al., 2022). For this reason, a maximum crown clade definition is used for the clade. Additional sequence data should be obtained to clarify the phylogenetic position of *S. graveolens*. Taxonomic treatments are available for *Solanum* section *Pachyphylla* (Dunal) Dunal (as the former genus *Cyphomandra*, Bohs, 1994) and *Solanum* section *Cyphomandropsis* (Bohs, 2001). Phylogenies for **/***Cyphomandra* include Bohs (2007), Särkinen & al. (2013), and Gagnon & al. (2022).

> **/*Geminata*** D. Don 1838 [S. Knapp & L.L. Giacomin] converted clade name. **Regnum number**: 1137.—**Definition:** The minimum crown clade containing *Solanum nudum* Dunal 1816 (*Solanum aphyodendron* S. Knapp 1985), *Solanum pseudocapsicum* Linnaeus 1753, *Solanum havanense* Jacquin 1794 and *Solanum reductum* C.V. Morton 1976.

**Primary reference phylogeny:** This paper (Fig. 1B). See also this paper (Fig. S1); Bohs (2005: Fig. 1); Gagnon & al. (2022: Fig. S12). **Etymology**: The name comes from the Latin “geminus” (twin), in reference to the paired leaf arrangement common in this clade. **Composition:** Includes 156 species of small shrubs to small trees primarily native to the American tropics, with a single species, *Solanum spirale* Roxb., native to Asia and Australia. Most species were previously classified as *Solanum* section *Geminata* (G.Don) Walp. **Diagnostic apomorphies**: A combined set of characters serves for recognition of /*Geminata*: absence of prickles, plants glabrous or with unbranched to dendritic trichomes and paired and often unequal leaves and leaf-opposed inflorescences in most species. **Synonyms:** None. **Comments:** This group of species was first described at subsectional rank as *Solanum* subsection *Geminata* G.Don. The species comprising the traditional *Solanum* section *Geminata* have also been referred to as section *Leiodendra* (Dunal) D’Arcy (e.g., D’Arcy, 1972, but see Knapp, 1983). Global taxonomic treatments are provided in Knapp (2002b, 2008) and for Brazil Knapp & al. (2015), and a phylogenetic treatment is provided by Tovar & al. (submitted to this volume). The type species of *Solanum* section *Geminata* (*Solanum nudum*) was not sampled for the primary reference phylogeny but was included in the more densely sampled phylogeny of Gagnon & al. (2022), where it belonged to a clade including *S. pseudocapsicum*. Tovar & al. (submitted to this volume) resolve *S. nudum* and *S. aphyodendron* in the same internal clade; both *S. aphyodendron* and *S. pseudocapsicum* were sampled in the primary reference phylogeny. *Solanum trachytrichium* Bitter (section *Silicosolanum* Bitter) and *Solanum apiahyense* Witasek were originally thought to belong to **/***Geminata* (Knapp, 2002b; Knapp & al., 2015) but recent phylogenetic studies (Gagnon & al., 2022) indicate that they are members of or more closely related to **/***Brevantherum*.

> **/*Aculeigerum*** Seithe 1962 [S. Knapp & L. Bohs] converted clade name. **Regnum number**: 1138.—**Definition:** The minimum crown clade containing *Solanum wendlandii* J.D. Hooker 1887 and *Solanum alternatopinnatum* Steudel 1841.

**Primary reference phylogeny:** This paper (Fig. 1B). See also this paper (Fig. S1); Gagnon & al. (2022: Fig. S12). **Etymology**: Derived from *Solanum* section *Aculeigerum* Seithe, and refers to the abundant prickles on all parts of the plants. **Composition:** Nine species of woody or semi-woody vines from the American tropics. **Diagnostic apomorphies**: Vines with recurved prickles lacking stellate hairs. **Synonyms:** None. **Comments:** Bohs (2005) showed a relationship between the taxa treated here as /*Aculeigerum* and *Solanum allophyllum* Standl. (section *Allophyllum* (A.Child) Bohs) and identified a group that has been informally referred to as “Wendlandii-Allophyllum” (e.g., Gagnon & al., 2022; Hilgenhof & al., 2023). *Solanum allophyllum* and morphologically similar taxa group together in Gagnon & al. (2022) and are sister to /*Nemorense*. But in our analysis here (see Fig. S2) the placement of *S. allophyllum* is not resolved, its position varies considerably in different analyses and further work is needed. The members of /*Aculeigerum* were treated monographically by Clark & al. (2015). For current clade composition see Hilgenhof & al. (2023).

> **/*Nemorense*** Child 1983 [L. Bohs] converted clade name. **Regnum number**: 1139.—**Definition:** The minimum crown clade containing *Solanum nemorense* Dunal 1813 and *Solanum reptans* Bunbury 1841.

**Primary reference phylogeny:** This paper (Fig. 1B). See also this paper (Fig. S1). **Etymology**: Derived from the specific epithet of the most widespread species in the group, *S. nemorense* (Child, 1983). **Composition:** Four species of shrubs and woody vines from the American tropics. **Diagnostic apomorphies**: Plants with prickles but lacking stellate hairs. **Synonyms:** none. **Comments:** A taxonomic revision of *Solanum* section *Nemorense* A.Child was published by Child (1983), who included several taxa not now considered related to *S. nemorense* in his circumscription of the section.

> **/*Leptostemonum*** Dunal 1852 [S. Knapp & T. Särkinen] converted clade name. **Regnum number**: 1140.—**Definition:** The minimum crown clade containing *Solanum polygamum* Vahl 1794, *Solanum mammosum* Linnaeus 1753, *Solanum macrocarpon* Linnaeus 1753, *Solanum melongena* Linnaeus 1753, and *Solanum vansittartense* C.A. Gardner 1923.

**Primary reference phylogeny:** This paper (Fig. 1B). See also this paper (Fig. S1); Bohs (2005: Fig. 1), Gagnon & al. (2022: Fig. S12). **Etymology**: From the Greek “leptos” (thin or slender) and “stemon” (stamen or thread), with reference to the strongly tapered anthers typically found in this group. **Composition:** Approximately 600 species of herbs, shrubs, vines, and small trees with a worldwide distribution. **Diagnostic apomorphies**: Most species have prickles, stellate hairs, and tapered anthers. **Synonyms:** *Solanum* subgenus *Leptostemonum* Bitter. **Comments:** This is the largest, most widely distributed, and most complex major clade in *Solanum* and comprises almost half the species diversity of the genus. It was first recognized at the sectional level by Dunal (1852). D’Arcy (1972) lists 46 different sectional and series names as belonging to subgenus *Leptostemonum*. Taxonomic overviews include those of Whalen (1984) and Nee (1999; New World species). Large-scale phylogenies include Stern & al. (2011; Americas), Vorontsova & al. (2013; Africa), Aubriot & al. (2016; tropical Asia), as well as more focused studies on smaller groups such as the eggplant and relatives (e.g., Aubriot & al., 2018) here recognised as **/***Melongena*. A large number of distinct clades are recognized in **/***Leptostemonum*, particularly in the American tropics; these will be treated under the PhyloCode in a separate publication (for a list and preliminary definitions of these, see Hilgenhof & al. 2023 and Särkinen & al., in prep).

> **/*Melongena*** Miller 1768 [S. Knapp & T. Särkinen] converted clade name. **Regnum number**: 1141. **Definition**: The minimum crown clade including *Solanum melongena* Linnaeus 1753 and *Solanum usambarense* Bitter & Dammer 1923.

**Primary reference phylogeny:** This paper (Fig. 1B). See also this paper (Fig. S1); Aubriot & al. (2018: Fig. 1); Gagnon & al. (2022: Fig. S12). **Etymology:** From the name applied to the cultivated eggplant *Solanum melongena* L. by Philip Miller (1768) to distinguish it at the generic level. **Composition:** Thirteen species of shrubs (one vine) mostly from Africa; *S. melongena* is widely cultivated, *S. incanum* L. is native from northern Africa to Pakistan, *S. insanum* L. is native to tropical Asia (see Vorontsova & Knapp, 2016; Ranil & al., 2016; Aubriot & Knapp, 2022). **Diagnostic apomorphies:** None. **Synonyms:** None. **Comments:** *Solanum melongena* and its closest relatives (see Knapp & al., 2013) have long been recognised as allied. Molecular phylogenies with increased sampling (Aubriot & al., 2016, 2018) expanded the membership of the clade to include the East African species *S. lanzae* J.-P.Lebrun & Stork, *S. agneworium* Vorontsova, and *S. usambarense*. Although support for this more inclusive clade is relatively weak in our reference phylogeny, inclusion of these three additional species is strongly supported in full plastome phylogenies (Aubriot & al., 2018).

> **/*Hemigeacanthos*** E. Gagnon & S. Knapp, new clade name. **Regnum number**: 1142.— **Definition:** The minimum crown clade containing *Solanum aculeastrum* Dunal 1852, *Solanum vansittartense* C.A. Gardner 1923, *Solanum mahoriense* D’Arcy & Rakotozafy 1986, *Solanum pubescens* Willd. 1814, and *Solanum anguivi* Lamarck 1794.

**Primary reference phylogeny:** This paper (Fig. 1B). See also this paper (Fig. S1); Gagnon & al. (2022: Fig. S12). **Etymology**: The name is derived from the Greek “hemi” (half), and “gaea/gea” (or “gaia”, the Earth) and refers to the distribution of the members of the clade in primarily the Eastern Hemisphere and to the possession of prickles (“acanthos”). **Composition:** With ca. 350 species, this clade is the largest within /*Leptostemonum* and includes all spiny solanums (with a few exceptions for species belonging to otherwise American clades, see Aubriot & al., 2016; Hilgenhof & al. 2023) whose native distribution occurs outside of the Americas. **Diagnostic apomorphies**: None except geography. **Synonyms:** None. **Comments:** This large group corresponds to the “Old World spiny solanums” (Stern & al., 2011; Vorontsova & al., 2013; Echeverría-Londoño & al., 2020) and to the “Eastern Hemisphere spiny solanums” of Gagnon & al. (2022) and Hilgenhof & al. (2023). The members of this large monophyletic group have been defined as distinct groups (e.g., Bitter, 1917, 1921, 1923; D’Arcy, 1972) or have been included in spiny solanum groups largely composed of American taxa (e.g., Whalen, 1984). D’Arcy (1972) listed 32 groups at series or sectional level that were typified by taxa included in /*Hemigeacanthos*. The history of classification of members of the group can be found for Australian and New Guinea species in Symon (1981, 1985), for Pacific species (McClelland, 2012), for African species (Vorontsova & Knapp, 2016), and for tropical Asian species (Aubriot & Knapp, 2022).

