## Supplementary figures and images for "A new phylogeny and phylogenetic classification for Solanaceae"

### Supplementary figure S1

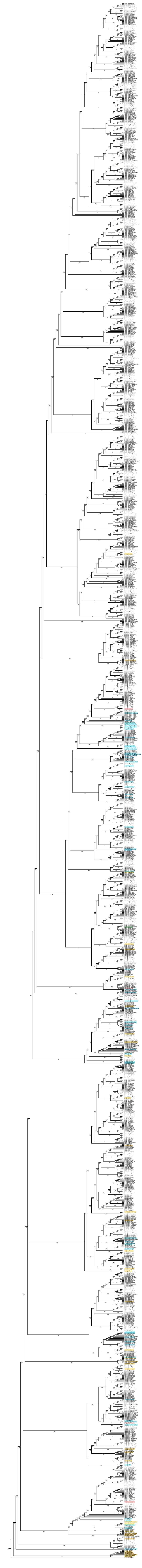

### Supplementary figure S2

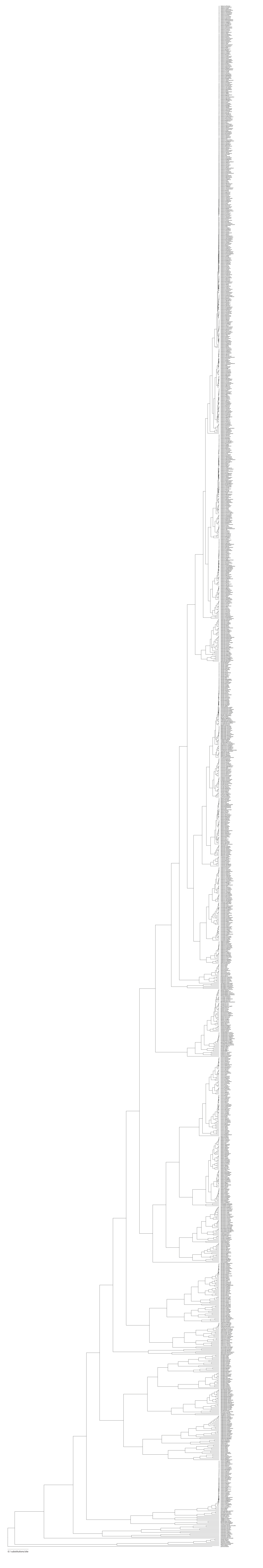

### Supplementary figure S4

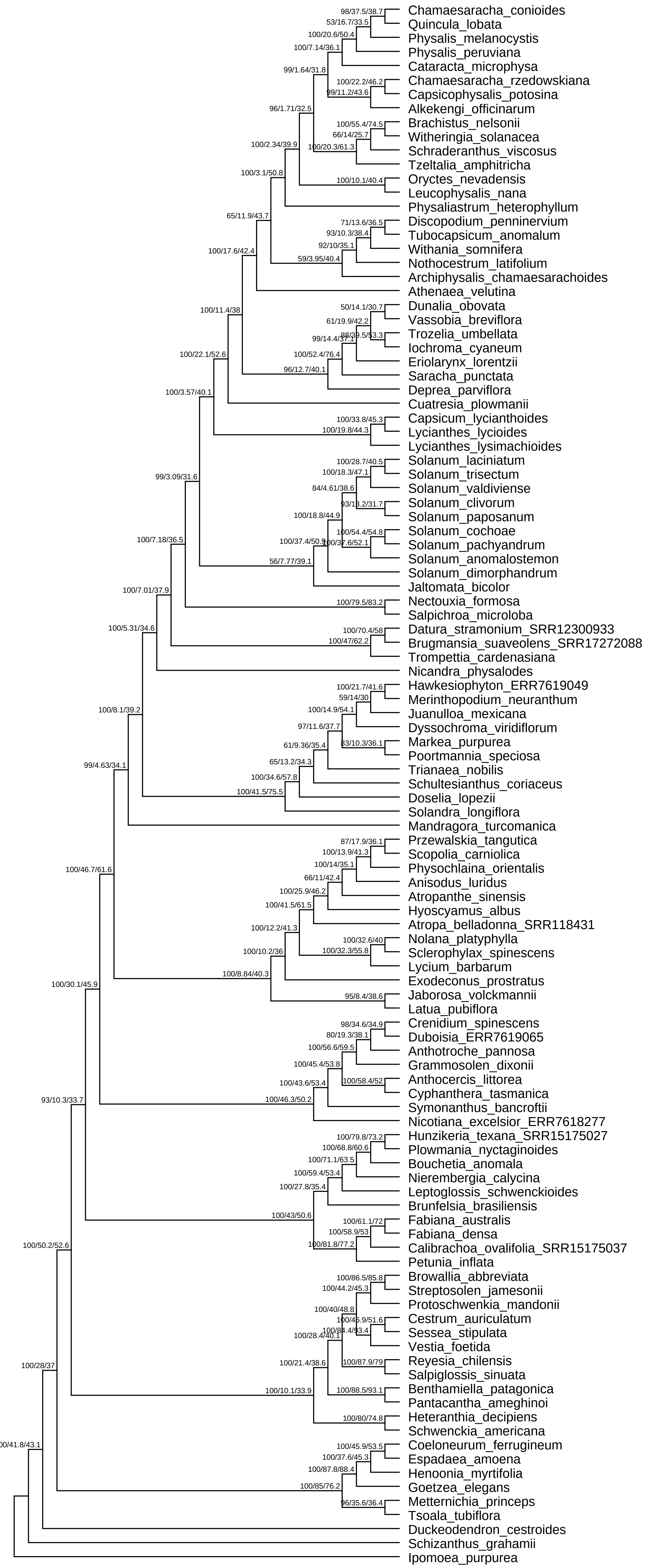

### Supplementary figure S5

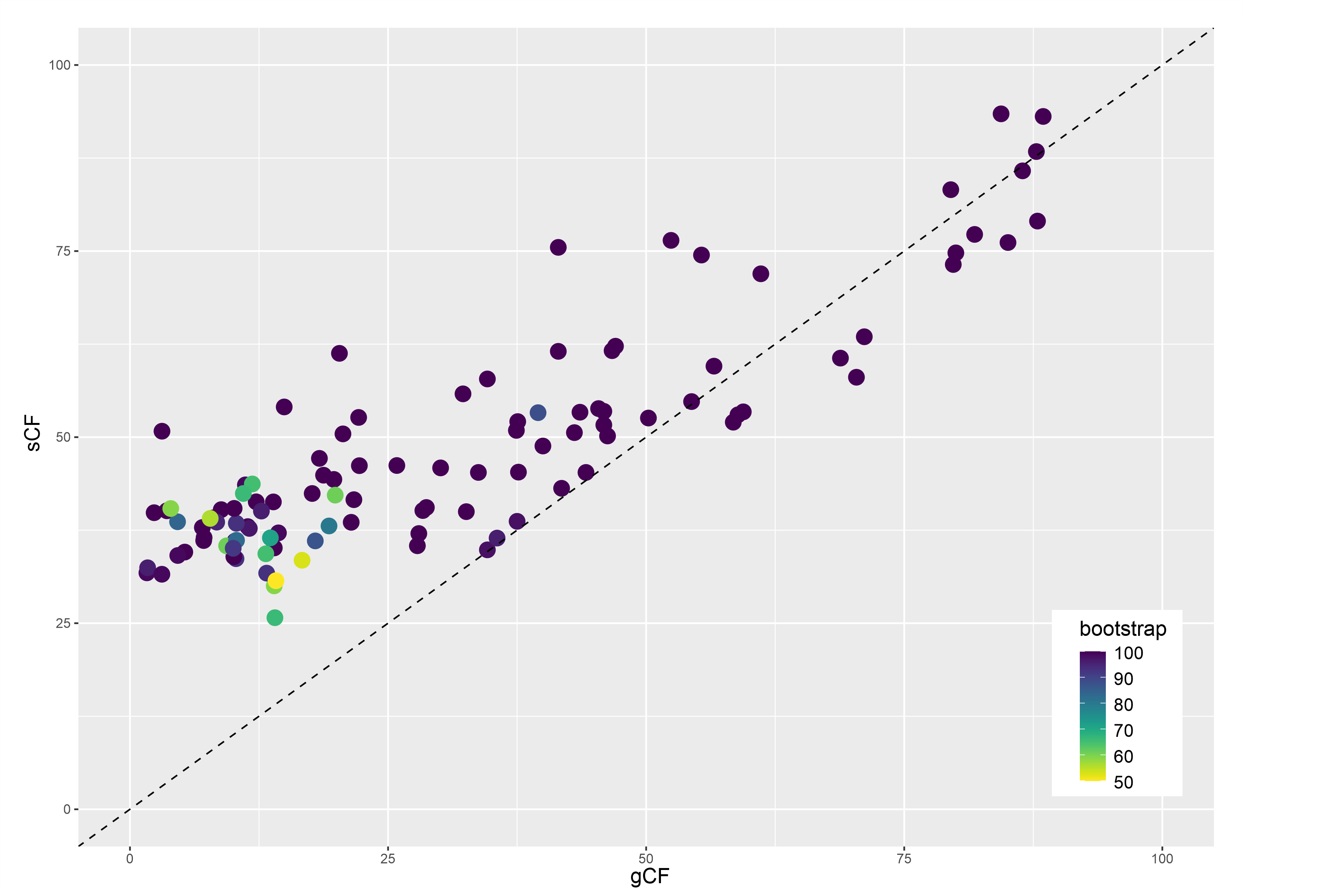
