## Supplementary table S2 for "A new phylogeny and phylogenetic classification for Solanaceae"

**Table S2.** Rogue taxa removed after two rounds of RogueNaRok cleaning (Aberer & al., 2013) based on trees generated from fast RAxML bootstrap analyses, using RAxML-VI-HPC v.8 (Stamatakis, 2014). Where the name in the original dataset is the same as the accepted name, the second column is left blank.

| **Name in the original dataset** | **Accepted name (if different)** |
| --- | --- |
| *Athenaea fasciculata* (Vell.) I.M.C.Rodrigues & Stehmann |  |
| *Brunfelsia macroloba* Urb. |  |
| *Capsicum buforum* Hunz. |  |
| *Capsicum ciliatum* (Kunth) Kuntze |  |
| *Capsicum cornutum* (Hiern) Hunz. |  |
| *Capsicum scolnikianum* Hunz. |  |
| *Cestrum acutifolium* Alain |  |
| *Cestrum bracteatum* Link & Otto |  |
| *Cestrum latifolium* Lam. |  |
| *Cestrum nocturnum* L. |  |
| *Cuatresia hunzikeriana* (Benítez & M.Martínez) N.W.Sawyer |  |
| *Datura quercifolia* Kunth |  |
| *Deprea hawkesii* (Hunz.) Deanna |  |
| *Deprea purpurea* (S.Leiva) Barboza & S.Leiva |  |
| *Hyoscyamus tibesticus* Maire |  |
| *Iochroma ellipticum* (Hook.f.) Hunz. |  |
| *Iochroma peruvianum* (Dunal) J.F.Macbr. |  |
| *Iochroma salpoanum* S.Leiva & K.Lezama |  |
| *Jaborosa volkmannii* (Phil.) Reiche |  |
| *Jaltomata cajamarca* Mione |  |
| *Lycianthes kaernbachii* (Lauterb. & K.Schum.) Bitter |  |
| *Lycium arenicola* MIers |  |
| *Lycium edgeworthii* Dunal | = *Lycium depressum* Stocks |
| *Lycium prunus-spinosa* Dunal | = *Lycium cinereum* Thunb. |
| *Nicotiana debneyi* Domin | = *Nicotiana forsteri* Roem. & Schult. |
| *Nicotiana didepta* | [artificial hybrid, name not validly published] |
| *Nicotiana kawakamii* Y.Ohashi |  |
| *Nolana albescens* (Phil.) I.M.Johnst. |  |
| *Nolana glauca* (I.M.Johnst.) I.M.Johnst. |  |
| *Nolana mollis* (Phil.) I.M.Johnst. |  |
| *Nolana platyphylla* (I.M.Johnst.) I.M.Johnst. |  |
| *Nolana reichei* M.O.Dillon & Arancio |  |
| *Nolana revoluta* Ruiz & Pav. |  |
| *Nolana spathulata* Ruiz & Pav. |  |
| *Nolana spergularioides* Ferreyra |  |
| *Nolana tovariana* Ferreyra |  |
| *Petunia guarapuavensis* T.Ando & Hashim. | = *Petunia scheideana* L.B.Sm. & Downs |
| *Physaliastrum japonicum* (Franch. & Sav.) Honda |  |
| *Physalis fuscomaculata* Dunal | = *Physalis viscosa* L. |
| *Physalis mendocina* Phil. | = *Physalis viscosa* L. |
| *Physalis subilsiana* J.M.Toledo |  |
| *Salpichroa microloba* Keel |  |
| *Schizanthus tricolor* Grau & E.Gronbach |  |
| *Scopolia parviflora* (Dunn) Nakai |  |
| *Sessea corymbiflora* R.Taylor & R.Phillips |  |
| *Solanum achacachense* Cárdenas | = *Solanum candolleanum* Berthault |
| *Solanum ambosinum* Ochoa | = *Solanum candolleanum* Berthault |
| *Solanum andreanum* Baker |  |
| *Solanum arcanum* Peralta |  |
| *Solanum batoides* D'Arcy & Rakot. |  |
| *Solanum bonariense* L. |  |
| *Solanum burtt-davyi* Dunkley | = *Solanum richardii* Dunal |
| *Solanum callium* R.J.F.Hend. | = *Solanum spirale* Roxb. |
| *Solanum capsicibaccatum* Cárdenas |  |
| *Solanum castaneum* Carvalho |  |
| *Solanum cataphractum* Benth. |  |
| *Solanum clavatum* Rusby |  |
| *Solanum delagoense* Dunal (= *Solanum campylacanthum* A.Rich) | probably *Solanum aligerum* Schltdl. |
| *Solanum didymum* Dunal |  |
| *Solanum drymophilum* O.E.Schulz | = *Solanum ensifolium* Dunal |
| *Solanum eremophilum* F.Muell. |  |
| *Solanum florulentum* Bitter | = *Solanum tarderemotum* Bitter |
| *Solanum hazenii* Britton |  |
| *Solanum hirsutissimum* Standl. | = *Solanum pectinatum* Dunal |
| *Solanum hispidum* Pers. |  |
| *Solanum humboldtii* Willd. | = *Solanum lycopersicum* L. |
| *Solanum humile* Lam. |  |
| *Solanum imamense* Dunal |  |
| *Solanum ipomoeoides* Chodat & Hassl. | = *Solanum uncinellum* Lindl. |
| *Solanum kurzii* Prain | = *Solanum violaceum* Ortega |
| *Solanum kwebense* C.H. Wright | = *Solanum tettense* Klotzsch |
| *Solanum latifolium* Poir. | = *Solanum rigidum* Lam. |
| *Solanum macranthum* Dunal | = *Solanum crinitum* Lam. |
| *Solanum manaense* Lemée | = *Solanum aturense* Dunal |
| *Solanum metarsium* C.V.Morton | = *Solanum sinuatirecurvum* Bitter |
| *Solanum mochiquense* Ochoa |  |
| *Solanum nitidum* Ruiz & Pav. |  |
| *Solanum obliquum* Ruiz & Pav. |  |
| *Solanum ovalifolium* Dunal |  |
| *Solanum panduriforme* Dunal | = *Solanum campylacanthum* A.Rich. |
| *Solanum parishii* A.Heller | = *Solanum umbelliferum* Eschsch. |
| *Solanum pronum* Thulin | = *Solanum cordatum* Forssk. |
| *Solanum pseudoquina* A.St.Hil. |  |
| *Solanum ptychanthum* Dunal | probably *Solanum emulans* Raf. |
| *Solanum racemosum* Jacq. | = *Solanum bahamense* L. |
| *Solanum ramonense* C.V.Morton & Standl. |  |
| *Solanum sanitwongsei* Craib | = *Solanum violaceum* Ortega |
| *Solanum scabrifolium* Ochoa |  |
| *Solanum sibundoyense* (Bohs) Bohs |  |
| *Solanum stenophyllidium* Bitter |  |
| *Solanum stipuloideum* Rusby |  |
| *Solanum subinerme* Jacq. |  |
| *Solanum tequilense* A.Gray | = *Solanum candidum* Lindl. |
| *Solanum trachytrichium* Bitter |  |
| *Solanum tridynamum* Dunal | = *Solanum houstonii* Martyn |
| *Solanum uleanum* Bitter |  |
| *Solanum uporo* Dunal | = *Solanum viride* Spreng. |
| *Trianaea naeka* S.Knapp |  |
| *Withania frutescens* (L.) Pauquy |  |
